# Ist2 promotes lipid transfer by Osh6 via its membrane tethering and lipid scramblase activities

**DOI:** 10.1101/2025.05.06.652365

**Authors:** Alicia Fabbre, Camille Syska, Heitor Gobbi Sebinelli, Anna Cardinal, Maud Magdeleine, Noha Al-Qatabi, Juan Martín D’Ambrosio, Cédric Montigny, Guillaume Lenoir, Alenka Čopič, Guillaume Drin

## Abstract

Lipid transfer proteins (LTPs) are required for the uneven distribution of lipids between cellular membranes, which is essential for many cell functions. In yeast, Osh6 is an LTP that exchanges phosphatidylserine (PS) with phosphatidylinositol 4-phosphate (PI(4)P) between the endoplasmic reticulum (ER) and the plasma membrane (PM), promoting the enrichment of PS in the PM. Here, we address why, to function optimally, Osh6 must bind to Ist2, an ER-resident lipid scramblase able to connect the ER to the PM via an intrinsically disordered region (IDR). We determined in vitro that Osh6 binds to the Ist2 IDR with micromolar affinity, whether empty or bound to its lipid ligands. Moreover, we found that Osh6 efficiently transfers PS at ER-PM contact sites if the Ist2 IDR has a minimal length and its binding site in the IDR is sufficiently removed from the ER surface. Next, we reconstituted the Osh6:Ist2 complex within artificial ER-PM contact sites and demonstrated that the association of Osh6 with Ist2 allows for a fast and directed PS flux between the connected membranes. We identified the Ist2 binding site on the Osh6 surface by validating structural models using our functional assays. Finally, we found that the Osh6-mediated PS transfer can be coupled to the PS scramblase activity of Ist2. These data unveil new functional partnerships between an LTP and a membrane tethering/scramblase protein and point to the general advantage of localizing these processes to membrane contact sites to ensure their efficiency.

## Introduction

Lipids are unevenly distributed within eukaryotic cells, which is essential for many cellular processes, like signal transduction or vesicular trafficking, as well as for cellular architecture (1–3). This is especially true for phosphatidylserine (PS): like most other glycerophospholipids, PS is synthesized at the endoplasmic reticulum (ER), but its concentration in the ER membrane is low, whereas it is highly enriched (up to five-fold) at the plasma membrane (PM) and the endosomes (4). PS contributes to the negative charge of the PM and is critical for the recruitment and activation of various signaling proteins (5). How PS becomes selectively enriched at the PM was largely unknown until the discovery that in yeast cells, two members of the oxysterol-binding protein(OSBP)-related protein (ORP) family (6), Osh6 and its close homolog Osh7, are lipid transfer proteins (LTPs) that carry PS from the ER to the PM (7). We have further shown that Osh6 exchanges PS for phosphatidylinositol 4-phosphate (PI(4)P), carrying PS from the ER to the PM coupled with the downhill transport of PI(4)P from the PM to the ER (8). The PI(4)P concentration gradient is maintained thanks to the PI-4 kinase Stt4, which generates PI(4)P at the PM in an ATP-dependent manner (9), and to the phosphatase Sac1, which degrades PI(4)P at the ER (10, 11).

In parallel, it has been reported that the human ORP5 and ORP8 function as PS/PI(4)P exchangers at ER-PM contact sites (12, 13), where the distance between the ER and the PM is less than 20 nanometers (14). These proteins have a more elaborate organisation than Osh6: in addition to a lipid transfer domain (i.e., OSBP-related domain, ORD) closely related to that of Osh6, they contain a C-terminal transmembrane domain to anchor them to the ER and an N-terminal phosphoinositide-binding PH domain to target the PM. This molecular organization allows them to engage in ER-PM contact sites and efficiently transfer lipids. However, in yeast cells, Osh6 and Osh7 can likewise be observed at the cortical ER, representing sites of close apposition between the ER and the PM (15–17), although they comprise only a single ORD and have no tethering domains/motifs. We and others uncovered that the cortical localization of Osh6 depends on its interaction with Ist2 (16, 17), one of the main proteins that scaffold ER-PM contact sites in yeast (18–21).

Ist2 consists of an N-terminal transmembrane (TM) domain embedded in the ER membrane (590 aa), followed by an extended, disordered cytosolic region of ∼360 amino acids (22). This intrinsically disordered region (IDR) permits Ist2 to interact with the surface of the PM via a short polybasic motif at the C-terminus that can bind specifically to PI(4,5)P_2_, a signaling phosphoinositide enriched in this membrane (23, 24). Deletion of Ist2 leads to a decrease in the ER area associated with the PM, but most of these contact sites are still maintained by other ER-PM tethering proteins (20, 21). By contrast, the function of the ER-embedded domain of Ist2 has long remained enigmatic. It comprises 10 predicted TM helices and bears homology to the TMEM16 protein family, whose members have been shown to function as phospholipid scramblases, facilitating lipid flip-flop across the bilayer structure of the membrane (25, 26). Although an initial study did not uncover any scrambling activity in proteoliposomes reconstituted with Ist2 (27), two recent studies demonstrated that the ER domain of Ist2 possesses Ca^2+^-independent phospholipid scramblase activity that influences processes at the ER, like vesicular transport and lipid droplet biogenesis (28, 29). Furthermore, cryo-EM analysis confirmed that the transmembrane domain of Ist2 structurally resembles TMEM16 proteins (28, 29).

Osh6 interacts with Ist2 by binding to a short motif in its disordered cytosolic tail and transfers PS at ER-PM contact sites (16, 17). Mutations of Osh6 or Ist2 that prevent this interaction result in cytosolic localization of Osh6 and a substantial defect in the PS transfer to the PM (16). However, why Osh6 must form a complex with Ist2 to maintain efficient PS transfer remains enigmatic. Although membrane contact sites are considered essential hubs for inter-organelle lipid transfer because many LTPs localize there (30), the advantage of this organization for the cell has been highly debated (31–34). Several possibilities have been proposed, which are not mutually exclusive.

First, it has been proposed that the localization of LTPs to membrane contact sites enables faster lipid transport. The idea is that a short distance between membranes might reduce the time needed for an LTP to move from one membrane to the other during a transport cycle; however, this hypothesis has not been addressed experimentally and remains disputed (33, 34). A second hypothesis is that the confinement of LTPs to contact sites guarantees a high degree of accuracy in transport, as it would prevent mistaken delivery of their lipid ligands to another organelle (31, 32). A third possible benefit is that the colocalization of LTPs to membrane contact sites enables or favours coordination between different lipid transport processes. In this respect, the interaction between Osh6 and Ist2 is particularly intriguing in light of the newly established lipid scrambling activity of Ist2. In eukaryotic cells, it is unclear whether PS is predominantly present in the cytosolic or luminal leaflet of the ER (35, 36) and thus easily accessible to a cytosolic protein. Recent structural data on human PS synthase 1 indicate that PS is synthesized in the luminal ER leaflet (37), highlighting the need for PS scrambling at the ER. More generally, an important question is whether lipid scramblases and LTPs can function together to ensure robust lipid fluxes between cellular membranes while maintaining their bilayer architecture. For instance, cellular and structural data suggest that rod-shaped LTP ATG2 works with two ER scramblases, TMEM41B and VMP1, and ATG9, a scramblase embedded in the nascent autophagosome to promote autophagosome expansion with phospholipids made in the ER (38–42). Yet, this hypothesis is only supported by a unique *in vitro* kinetic measurement suggesting that ATG9 enhances ATG2-mediated lipid transfer between membranes (43). Thus, there is a lack of data on whether a lipid scramblase activity can be biochemically coupled to an intermembrane lipid transfer process.

Here, we have analyzed the functional coupling between Osh6 and Ist2 by combining biochemical approaches, *in vitro* reconstitution assays, cellular observations, and structural modeling. We demonstrate that Osh6 can associate with the IDR of Ist2 with a micromolar affinity, independent of its loading with a lipid ligand. We test in cells how the Ist2 IDR contributes to PS transfer to the PM, and show that PS transport is delayed if the IDR is too short or if the Osh6-binding site is too close to the ER surface. Then, we recapitulate the capacity of Ist2 to connect ER- and PM-like membranes and to recruit Osh6 between these membranes. Using this reconstituted system, we demonstrate that the association of Osh6 with the Ist2 IDR promotes a fast and directed flux of PS between membranes connected by the IDR. We further support this observation by generating Osh6 mutants that are unable to interact with Ist2 and do not show increased PS transfer activity between tethered liposomes. Finally, by combining *in vitro* PS transfer assays with lipid scrambling assays using Ist2 proteoliposomes, we show that Ist2 can sustain the PS transfer activity of Osh6 thanks to its PS scrambling capacity. Collectively, these results provide the molecular basis for the functional coupling between an LTP and a dual membrane tether/lipid scramblase, which contributes to the proper distribution of PS in cells.

## Results

### Analysis of the interaction between Osh6 and the disordered region of Ist2

The C-terminal IDR of Ist2 contains about 360 aa. Evidence obtained in yeast cells and *in vitro* showed that Osh6 binds to a minimal motif in the region comprising residues 729 to 747 (16, 17). However, no characterization of how Osh6 interacts with the full-length Ist2 IDR exists. To achieve this aim, we first used GST pull-down assays. Three segments of the Ist2 IDR appended to an N-terminal GST were used as baits (**Fig. 1a**): a 727–776 segment encompassing the Osh6-binding motif of Ist2 as a positive control (17), a 590–768 segment in which this motif is at the C-terminus of the construct and potentially easily accessible, and a 590–946 segment corresponding to the entire disordered region of Ist2. We found that all three Ist2 segments fused with GST could recruit Osh6 to the beads and that the amount of Osh6 recruited was similar, regardless of the construct used (**Fig. 1b, c**). No binding was observed with GST alone or with an Ist2 construct with a truncated Osh6-binding motif (GST-Ist2[590–946]Δ727-749) (**Fig. S1**). The same experiments with ORD^ORP8^, ORD^Osh3,^ and Osh4 showed that these constructs were not recruited by the disordered region of Ist2, demonstrating the specificity of the Ist2:Osh6 interaction (**Fig. 1d, Fig. S2**). These data show that Osh6 can associate *in vitro* with the full-length Ist2 IDR by recognizing a binding motif site in its 727–776 segment.

**Figure 1.**
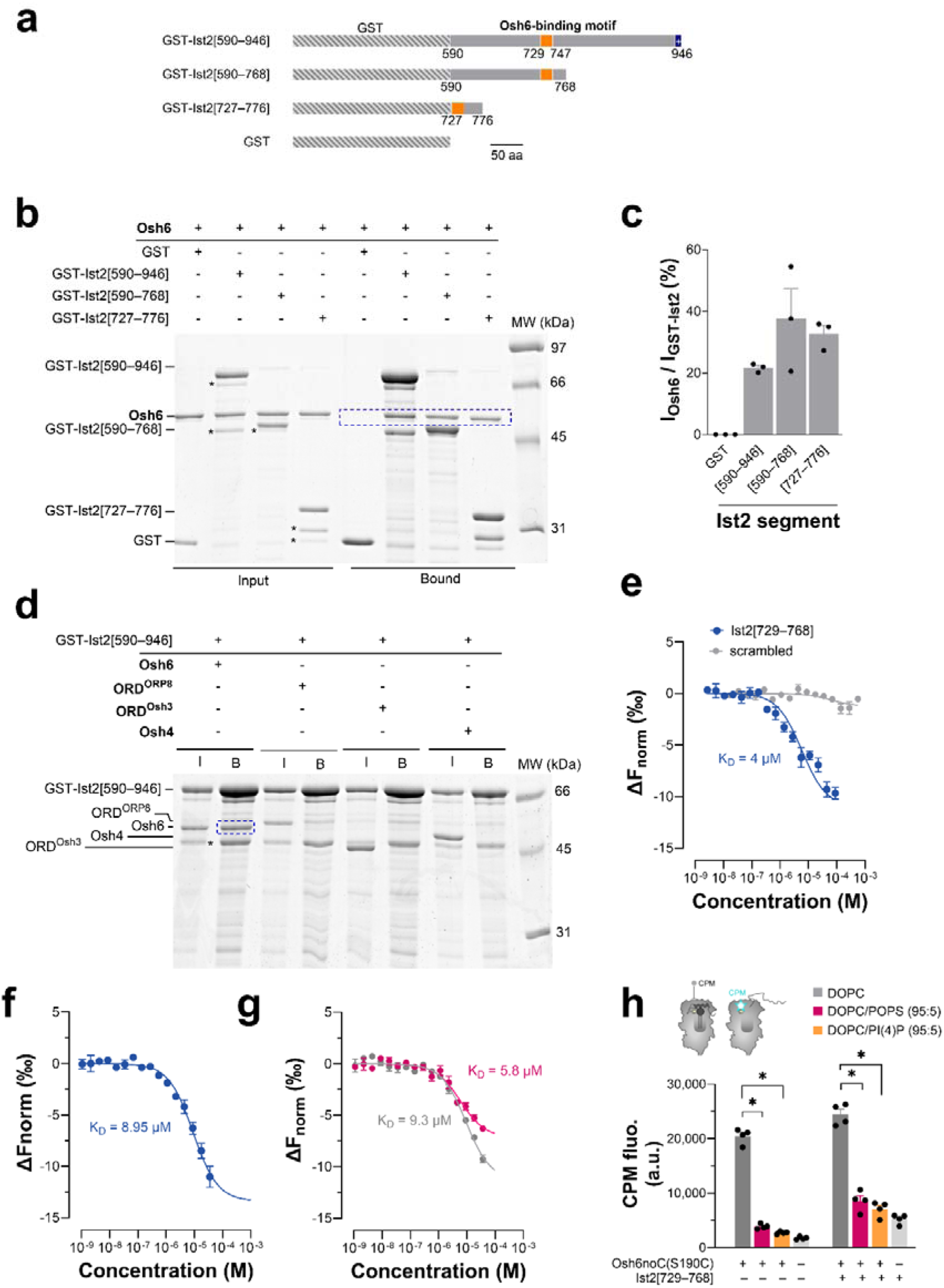
Interaction between Osh6 and the IDR of Ist2. **(a)** The capacity of Osh6 to bind to different segments of the Ist2 Intrinsically Disordered Region (IDR) has been examined by GST pull-down using GST-Ist2[590–946], GST-Ist2[590–768], and GST-Ist2[727–776] constructs. The orange segment corresponds to the Osh6-binding motif in the Ist2 IDR. **(b)** GST pull-down. GST-tagged segments of Ist2 IDR were immobilized on glutathione beads and incubated with Osh6. Input and bound fractions were analyzed on SDS-PAGE with SYPRO Orange staining. The stars indicate the main contaminants in GST-tagged construct preparation. As pointed out by the blue box, Osh6 interacts with the three GST-tagged segments of the Ist2 IDR. No interaction was seen with GST alone. **(c)** Quantification of the GST-pull down assays as shown in (b). Osh6 binds to the three GST-tagged segments of Ist2 IDR with almost the same efficiency. Data are represented as mean ± s.e.m. (n = 3) with single data points. **(d)** GST pull-down. GST-Ist2[590–946] was immobilized on beads and incubated with Osh6, ORD^ORP8^, ORD^Osh3^, or Osh4. Input (I) and bound (B) fractions were analyzed using SDS-PAGE. Contrary to Osh6, neither ORD^ORP8^, ORD^Osh3^, nor Osh4 can associate with the Ist2 IDR. **(e)** MST binding assay. Osh6 labeled with Alexa Fluor 647 C2-maleimide (Osh6-AF647, 20 nM) was incubated with sixteen different concentrations of Ist2[529–768] peptide in HBS-EP+ buffer at 25 °C for 10 min under shaking. Then, each sample was loaded in a capillary, and measurements were performed using a Monolith NT.115 instrument. An MST-on time of 5 s was set for the analysis, and baseline-corrected normalized fluorescence values (ΔF_norm_[‰]) were plotted against peptide concentration. Curves were fitted with a nonlinear regression model. A negative control experiment was performed with a peptide corresponding to a scrambled Ist2[529–768] sequence. No binding was observed (grey dot). Data are represented as mean ± s.e.m. (n = 3–4). **(f)** MST binding assay. Osh6-AF647 (20 nM) was mixed with different concentrations of Ist2[590– 936] construct in HBS-EP+ buffer at 25 °C for 10 min under shaking. Measurements and analysis were performed as in (e). Data are represented as mean ± s.e.m. (n = 3). **(g)** MST binding assay. Osh6-AF647 (20 nM), loaded or not with POPS, was mixed with different concentrations of Ist2[590–936] construct in HBS-EP+ buffer at 25 °C for 10 min under shaking. Measurements and analysis were performed as in (e). Data are represented as mean ± s.e.m. (n = 3). **(h)** Accessibility assay. CPM (4 µM) was mixed with 400 nM Osh6(noC/S190C) incubated with pure DOPC liposomes or liposomes doped with 5% POPS or brain PI(4)P, in the absence or presence of 40 µM Ist2[729–768]. Control experiments were done without Osh6(noC/S190C) and liposome. Intensity bars correspond to the fluorescence measured 25 min after adding CPM (n = 3–4).

Next, we measured Osh6’s affinity for the Ist2 IDR by microscale thermophoresis (MST). First, we produced a fluorescent version of Osh6 (AF647-Osh6) by coupling an Alexa Fluor 647-C2-maleimide dye to a unique cysteine (C262) lying at the protein surface (**Fig. S3a-c**, as shown later, we identified that the Ist2-binding site is away from this position; we were thus confident that the fluorescent dye would not perturb the capacity of Osh6 to interact with the Ist2 IDR). Then, we mixed AF647-Osh6 with an increasing concentration of a peptide corresponding to the Ist2[729–768] segment, encompassing the minimal Osh6 binding motif. MST traces indicated an interaction between Osh6 and this peptide, with a K_D_ of ∼4 µM (**Fig. 1e, Fig. S3d**). A control experiment with a peptide corresponding to a scrambled version of the Ist2[729–768] segment revealed no binding, validating the assay’s specificity.

Next, we produced the Ist2 IDR devoid of GST tag, with or without the C-terminal positively charged binding motif (Ist2[590–936] and Ist2[590–946], **Fig. S3e**) to perform MST measurements. However, we could only characterize Ist2[590–936], since Ist2[590–946] was not pure enough for spectroscopic analyses. We recorded the CD spectrum of Ist2[590–936], showing a minimum at λ = 198 nm (**Fig. S3f**), which indicated it was unfolded. Using dynamic light scattering (DLS), we found that Ist2[590–936], despite its smaller molecular weight compared to Osh6 (28 kDa vs. 51.6 kDa), has a higher hydrodynamic radius (5.3 versus 3.8 nm) (**Fig. S3g**), consistent with the notion that Ist2 IDR has a disordered structure. Finally, we measured the affinity of AF647-Osh6 for Ist2[590–936] by MST and obtained a K_D_ of ∼ 9 µM (**Fig. 1f**). These data indicated that Osh6 has no problem reaching its binding motif within the Ist2 IDR, with an affinity in the micromolar range.

Next, we examined how the Ist2 IDR binds to Osh6 in the apo or lipid-bound state. To this end, following an established protocol (44), we prepared AF647-Osh6 loaded with 1-palmitoyl-2-oleoyl-*sn*-glycero-3 phospho-L-serine (POPS) or PI(4)P, or in an empty form. We found that the affinity of AF647-Osh6 for Ist2[590–936] was similar (K_D_ = 5 vs. 9 µM, **Fig. 1g**), whether it was in the apo form or bound to POPS. With the PI(4)P-loaded form of AF647-Osh6, we obtained variations in the MST signal that were too low to provide a K_D_ value. However, we found using GST pull-down that Osh6 loaded with PI(4)P can bind to the Ist2 IDR as efficiently as an empty or PS-bound form of Osh6 (**Fig. S3h,i)**. Finally, we assessed whether Osh6 bound to the Ist2 IDR could bind to its lipid ligands. For this, we used an Osh6(noC/S190C) construct with a unique cysteine at position 190 that is solvent-exposed only when the lipid-binding pocket of the protein is empty, and the molecular lid that gates this pocket is open (44, 45). This construct was added to liposomes doped or not with 5 mol% POPS or PI(4)P in the presence or absence of the Ist2[729–768] peptide. Then, 7-diethylamino-3-(4′-maleimidylphenyl)-4-methylcoumarin (CPM), a molecule that becomes fluorescent only when forming a covalent bond with accessible S190C, was added to each sample. After a 25-min incubation, a high fluorescence signal was measured with Osh6(noC/S190C) mixed with PC liposomes, indicating that the protein remained open (**Fig. 1h**). In contrast, almost no fluorescence was recorded when the protein was incubated with PS or PI(4)P-containing liposomes, indicating that it remained mostly closed, as previously shown (44, 45). Identical results were obtained when the Ist2[729–768] peptide was incubated with Osh6. Altogether, these data established that the association of Osh6 with the Ist2 IDR and its loading with ligands are not mutually exclusive events, and that Ist2 IDR binding to Osh6 occurs away from the Osh6 lipid binding site.

### The influence of Ist2 IDR on Osh6-mediated PS transfer in cells

We next explored how the Ist2 IDR affects Osh6 localization and its PS transfer activity in a cellular context. For this, we constructed different variants of Ist2 with truncations of its IDR, which were tagged with blue fluorescent protein (BFP) at the N-terminus **(Fig. 2a)**. These variants were expressed in budding yeast from a low-copy plasmid and the native Ist2 promoter, as previously described (16). All variants contained an unperturbed C-terminal region, aa878–946, to allow efficient binding to the PM (24, 46). Deletion of this region resulted in improper localization of BFP-Ist2[1–877] in an *ist2Δ* background, likely implying protein aggregation **(Fig. S4a)**. Likewise, we could not express an N-terminal truncation mutant lacking the full transmembrane (TM) domain of Ist2, BFP-Ist2[590–946]. In this case, the fluorescent signal was very low and again sometimes observed in punctate structures, suggesting protein aggregation **(Fig. S4b)**. These results show that the IDR of Ist2 needs to be always tethered to two membranes and that the lack of tethering leads to its aggregation in cells. In agreement, the localization of BFP-Ist2[1–877] can be rescued by the addition of a CAAX motif that leads to protein prenylation and attachment of the IDR to the PM via this lipid anchor (BFP-Ist2[1–877]PM) **(Fig. S4c)**. This result further underlines the importance of Ist2-IDR membrane attachment for its localization in cells, whereas the C-terminal region itself can be deleted.

**Figure 2.**
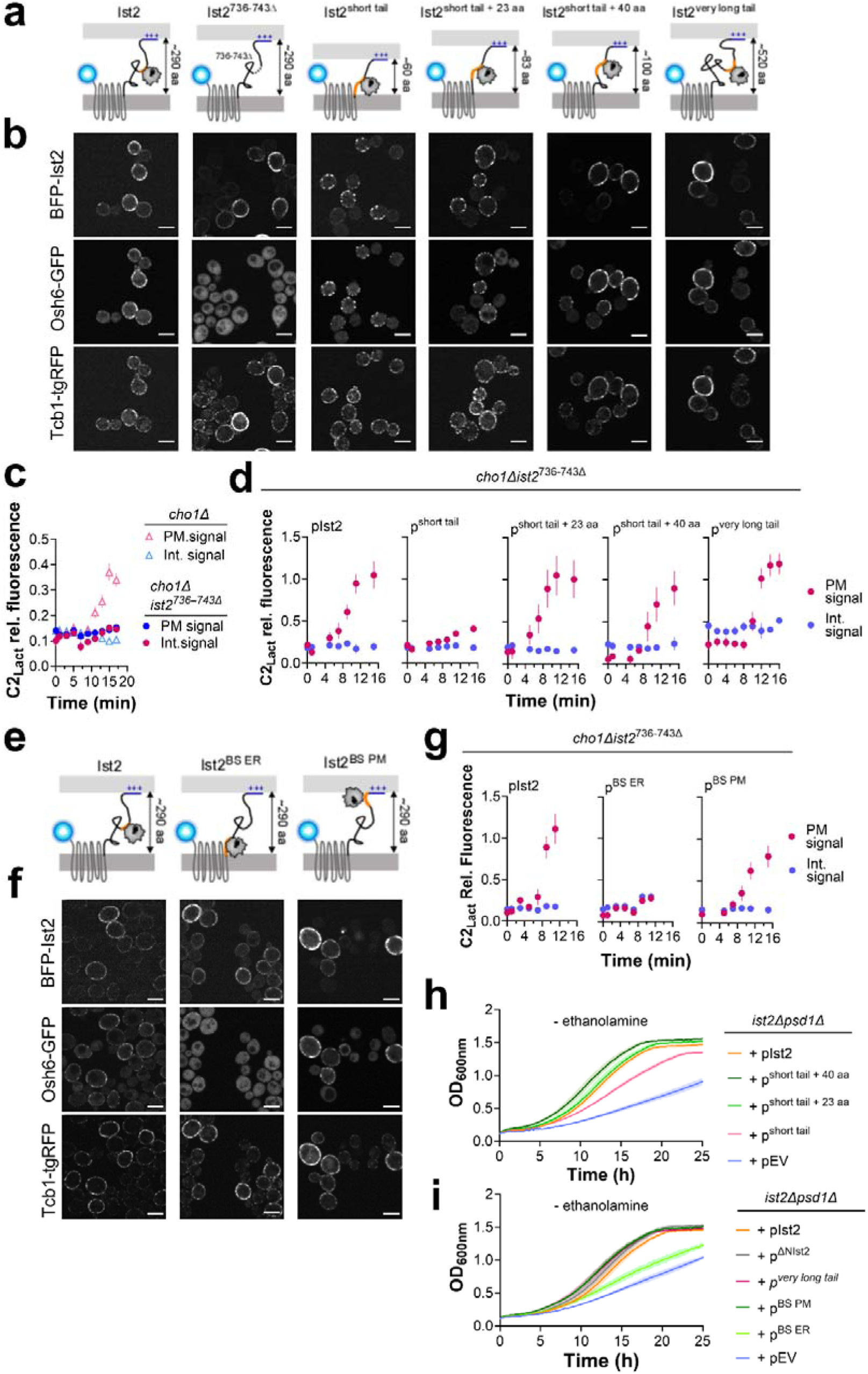
The influence of the Ist2 IDR on Osh6 localization and PS transport in cells. **(a)** Diagrams of Ist2 variants with truncations of the cytosolic tail, tagged N-terminally with BFP (light blue sphere). Length of the cytosolic tail between the ER-embedded TM region [1–590] and the extended PM-binding region ([878–946], dark blue), which interacts with the PM (light grey), is indicated. The Osh6 binding site is depicted in orange. **(b)** Localization of plasmid-encoded BFP-Ist2 variants in *ist2*Δ cells, and colocalization with genomically-tagged Osh6-GFP and Tcb1-tgRFP. Scale bar, 5 µm. **(c)** Evaluation of PS transport from the ER in *cho1Δ* cells. Localization of the PS probe C2_Lact_–GFP was evaluated simultaneously in two strains lacking PS, *cho1*Δ and *cho1*Δ *ist2^736–743^*^Δ^, over time every 2 min after the addition of lyso-PS, as described (16) (see also **Fig. S4d**). Intensity of C2_Lact_–GFP signal is plotted over time, comparing internal peaks (blue symbols) and peripheral (PM) peaks (red symbols, showing probe redistribution from internal (cytosolic, ER) towards peripheral (PM) peaks in *cho1*Δ (expressing Ist2-WT, triangles), but not in *cho1*Δ *ist2^736–743^*^Δ^ cells (round symbols), as previously shown (16). Data are mean ± s.e.m. (n = 10–15 cells) from one of two independent experiments. Scale bars, 5 μm. **(d)** Plasmids encoding BFP-Ist2 variants with different lengths of the cytosolic tail were introduced into *cho1*Δ *ist2^736–743^*^Δ^ and C2 –GFP localization was evaluated as in (c). Data are mean ± s.e.m. (n = 10 cells) from one of two independent experiments. **(e, f, g)** Diagram of Ist2 variants with Osh6-binding region [705–762] moved close to the ER (after [1– 589], Ist2^BS^ ^ER^), or close to the PM (before [878–946], Ist2^BS^ ^PM^) (see **Table S2** for details). Localization of these variants (f) was assessed as in (b), and their PS transport activity (f) was assessed as in (c). **(h, i)** Growth of *ist2Δ psd1Δ* cells with plasmids encoding different BFP-Ist2 variants, as indicated, was followed in minimal media without ethanolamine at 30 °C (control experiments with ethanolamine are shown in **Fig. S4e-g**). Data are mean ± s.e.m. from triplicate cultures showing one of two independent experiments.

When wild-type BFP-Ist2 was expressed in a strain with endogenously-tagged Osh6-GFP and lacking the chromosomal copy of *IST2* (*ist2Δ),* it showed a patchy distribution at the cell periphery typical for ER-PM contact sites, consistent with the previously-reported localization of Ist2 (19, 20, 22). Furthermore, it colocalized with another ER-PM contact site protein, Tcb1, which was endogenously labelled with tagRFP (tgRFP). Osh6-GFP co-localized with Ist2 and Tcb1 and showed some cytosolic signal (**Fig. 2b**, right panel). By contrast, when the binding site for Osh6 in Ist2 IDR was truncated (Ist2^736-743Δ^, (16)), Osh6-GFP was completely cytosolic, whereas the localization of Ist2 and Tcb1 was not affected. We then explored the effect of deleting other regions of Ist2 IDR, while preserving the full Osh6 binding region (aa704-761; (16)), on Ist2 and Osh6 localization. It was previously shown that the cytosolic portion of the C-terminal tail of Ist2 can be significantly truncated without a noticeable effect on Ist2 localization, until the truncation down to ∼60 aa, where Ist2 shows a punctate localization (16, 22). In agreement, our Ist2^short-tail^ mutant showed a more punctate localization at the cell periphery **(Fig. 2b)**. Addition of 23 or 40 aa between the TM domain and Osh6-binding region resulted in a normal Ist2 localization, and an increase in the length of the IDR by 230 aa (duplication of regions [590-704] and [763-877]) also did not significantly affect the Ist2 distribution nor its colocalization with Tcb1. In all cases, Osh6 colocalized with Ist2 at the cell periphery **(Fig. 2b)**.

We next used a cellular PS transfer assay to assess the influence of Ist2 IDR on the PS transfer activity of Osh6. This assay requires *cho1Δ* cells, which lack the sole PS synthase in yeast, Cho1, and therefore are devoid of PS (47). PS can be supplied exogenously in the form of lyso-PS, which is transported to the ER and acylated into PS, and whose transport back to the PM can be observed in real time using the fluorescent PS reporter C2_Lact_-GFP (7, 8, 48). As we have previously shown, C2_Lact_-GFP localizes to the PM within 15–30 min after lyso-PS addition in *cho1Δ IST2* cells, but remains intracellular and localized to the ER, in *cho1Δ ist2^736-743Δ^* cells (16) (**Fig. 2c, Fig. S4d).** We were not able to construct a *cho1Δ ist2Δ* strain due to synthetic lethality (28), therefore the effect of BFP-Ist2 IDR truncation mutants on PS transport was assessed in the *cho1Δ ist2^736-743Δ^* background **(Fig. 2d)**. In this background, only Ist2^short-tail^ showed a defect in PS transport, whereas the other truncation mutants, as well as Ist2^very^ ^long^ ^tail^ supported PS transport to a similar extent as wild-type Ist2. Furthermore, PS transport was also supported by the Ist2 prenylation mutant BFP-Ist2[1-877]PM, suggesting that the C-terminal PM-binding region of Ist2 is not essential for PS transport (result not shown).

We also asked how the position of the Osh6-binding region within the Ist2 IDR affects Osh6 localization and its PS transfer activity. This binding region is positioned roughly in the middle of the IDR. We constructed two Ist2 mutants, Ist2^BS^ ^ER^, where the binding site was moved next to the TM region, and Ist2^BS^ ^PM^, where the binding site was moved next to the C-terminal PM-binding region [878–946] **(Fig. 2e)**. Both Osh6 localization and PS transport were affected by the Ist2^BS^ ^ER^ mutant, but not by the Ist2^BS^ ^PM^ mutant **(Fig. 2f,g)**.

An issue with our PS transport assay is that Ist2 IDR mutants were tested in the presence of Ist2^736-743Δ^, which cannot bind to Osh6 but could compensate for the IDR *in trans*, especially given that Ist2 dimerizes via its TM domain (29). We therefore used an independent genetic approach to evaluate the effect of Ist2 IDR on Osh6 activity and cellular PS homeostasis, assessing the genetic interaction of *ist2* mutants with the PS-decarboxylase mutant *psd1Δ* (17). As previously shown, the growth of *ist2Δ psd1Δ* cells is strongly inhibited in synthetic media lacking ethanolamine, but is restored by the addition of ethanolamine **(Fig. 2h, Fig. S4e)**. Growth in this strain is rescued by the addition of WT Ist2 (pIst2), and by all Ist2 IDR mutants except for Ist2^short-tail^ **(Fig. 2h)** and by Ist2^BS^ ^ER^ **(Fig. 2i)**, or by ethanolamine **(Fig. S4e-g)**, in agreement with the results of the PS transport assay. Together, this analysis shows that the majority of Ist2 IDR is not required for Osh6 localization or PS transport activity in cells, at the sensitivity of our assays, as long as a minimal length is ensured and/or the Osh6-binding region is sufficiently removed from the ER surface.

### Osh6 is recruited by the Ist2 IDR attached to ER or PM mimetic membranes

To better understand why membrane tethering by Ist2 IDR is necessary for PS transfer by Osh6, we decided to reconstitute this process *in vitro*. To do so, we examined first whether it was possible to anchor the N-terminus of the disordered region of Ist2 to liposomes mimicking the ER membrane (**Fig. 3a**). We purified a _C_Ist2[590–946] construct (**Fig. S3e**) with an extra N-terminal cysteine, to covalently attach it to the surface of 1,2-dioleoyl-sn-glycero-3-phosphoethanolamine-N-[4-(p-maleimidophenyl)butyramide] (MPB-PE)-containing liposomes with an orientation comparable to that of the Ist2 IDR in a cellular context (it is worth noting that the Ist2 IDR does not contain endogenous cysteine) (**Fig. 3b**). Then, we prepared liposomes composed of DOPC to mimic the ER membrane, which is fluid and relatively neutral, and made of unsaturated lipids, additionally containing 0, 2, or 10 mol% MPB-PE. These liposomes (0.75 mM lipids) were mixed with DTT-free _C_Ist2[590–946] (0.75 µM). Then, by flotation assays, we quantified how much protein was associated with the liposomes collected at the top of a sucrose gradient after ultracentrifugation. As expected, we found that _C_Ist2[590–946] was weakly anchored to liposomes consisting only of DOPC (1.4 % of total protein compared to the condition without liposome) but 9-fold more to liposomes containing 2% MPB-PE and 27-fold more to liposomes containing 10% MPB-PE. These results indicate that the _C_Ist2[590–946] construct can be covalently attached via its N-terminal tip to liposomes containing 10% MPB-PE (**Fig. 3 b).** Next, we tested whether the Ist2 IDR could recruit Osh6 to the vicinity of the ER-like membrane. Liposomes, doped or not with 10% MPB-PE, were mixed with _C_Ist2[590–946], then treated with DTT to stop any subsequent coupling reaction with MPB-PE and mixed with Osh6. Flotation assays showed that Osh6 was only slightly recruited to DOPC-only liposomes in the presence of _C_Ist2[590–946] but 2-fold more if this construct was attached to MPB-PE-containing liposomes (**Fig. 3c**). These observations suggest that the Ist2 IDR, anchored via its N-terminus to the surface of neutral membranes, recruits Osh6.

**Figure 3.**
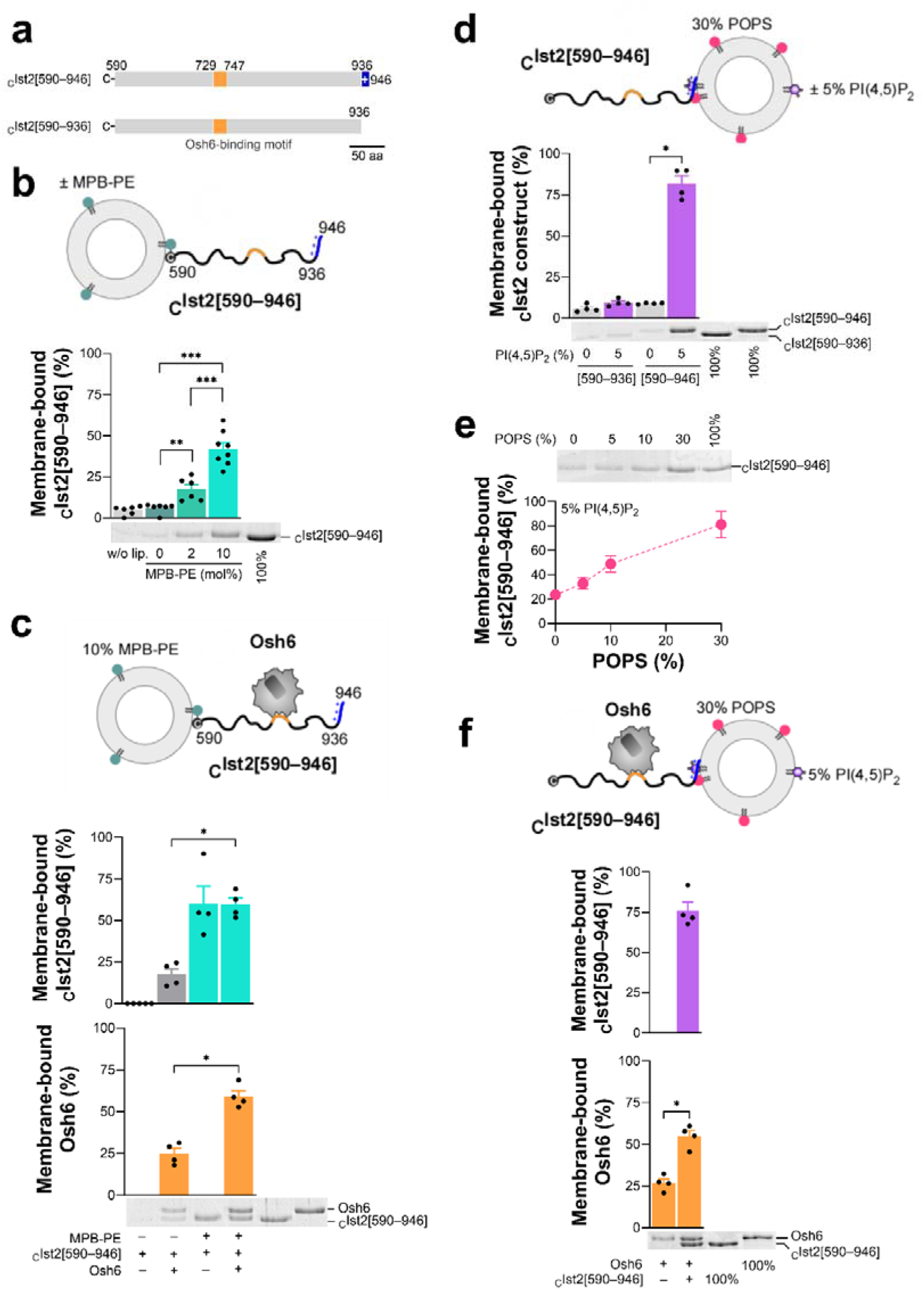
Osh6 is recruited by the Ist2 IDR, anchored by its N-terminal end to ER-mimicking liposomes or by its C-terminal end to PM-mimicking liposomes. **(a)** Schematic of the _C_Ist2[590–946] and _C_Ist2[590–936] constructs corresponding to the Ist2 IDR with or without the C-terminal basic motif, respectively; the two constructs contain an extra N-terminal cysteine. **(b)** Flotation assay. Liposomes made of DOPC, containing 0, 2, or 10% MPB-PE and 0.1% NBD-PE (750 µM total lipids) were mixed with DTT-free _C_Ist2[590–946] (0.75 µM) for 1 h in HK buffer at 25°C. A control experiment was made with protein but no liposomes. After centrifugation, the liposomes were collected at the top of sucrose cushions and analyzed by SDS-PAGE. The amount of membrane-bound protein was determined using the content of lane 5 (100% total) as a reference based on the SYPRO Orange intensity. Data are represented as mean ± s.e.m. (n = 6–8) with single data points. Unpaired Mann–Whitney U test (\**p* < 0.05, \*\**p* < 0.01). **(c)** Flotation assay. _C_Ist2[590–946] construct (0.75 µM) was mixed for 1 h with DOPC liposomes, doped or not with 10% MPB-PE and 0.1% NBD-PE. Then, DTT (2 mM) was added to stop the functionalization reaction, and the liposomes were mixed or not with 0.75 µM of Osh6 for 10 min. The amount of membrane-bound _C_Ist2[590–946] and Osh6 was determined using the content of lanes 5 and 6 (100% total), respectively. Data are represented as mean ± s.e.m. (n = 4) with single data points. Unpaired Mann–Whitney U test (\**p* < 0.05). **(d)** Flotation assay. Liposomes (750 µM lipids) composed of POPC/POPS (70:30) or POPC/POPS/PI(4,5)P_2_ (65:30:5), doped with 0.1% NBD-PE, were mixed with _C_Ist2[590–946] or _C_Ist2[590–936] (0.75 µM) for 10 min. Data are represented as mean ± s.e.m. (n = 4) with single data points. Unpaired Mann–Whitney U test (\**p* < 0.05, \*\**p* < 0.01). **(e)** Flotation assay. Liposomes (750 µM lipids) composed of POPC/PI(4,5)P_2_ (70:30) with increasing amounts of POPS (0, 5, 10 or 30% at the expense of POPC), doped with 0.1% NBD-PE, were mixed with _C_Ist2[590–946] (0.75 µM) for 10 min. Data are represented as mean ± s.e.m. (n = 4). **(f)** Flotation assay. Liposomes (750 µM lipids) composed of POPC/POPS/PI(4,5)P_2_/NBD-PE (65:30:5:0.1) were mixed with 0.75 µM of Osh6 in the presence or absence of an equivalent amount of _C_Ist2[590–946] for 10 min. Data are represented as mean ± s.e.m. (n = 4) with single data points. Unpaired Mann–Whitney U test (\**p* < 0.05).

In a second step, we examined whether the _C_Ist2[590–946] construct could interact with a membrane mimicking the inner leaflet of the PM via its C-terminal polybasic motif (23, 24). To this aim, we analyzed by flotation assays to what extent _C_Ist2[590–946] and the construct lacking the polybasic motif, _C_Ist2[590–936] (**Fig. S3e**), associated with liposomes composed of 70% 1-palmitoyl-2-oleoyl-*sn*-glycero-3-phosphocholine (POPC) and 30% POPS, doped or not with 5% PI(4,5)P_2_. We found that _C_Ist2[590–946] was firmly bound to liposomes containing PI(4,5)P_2_ but not to liposomes in which this lipid was lacking (**Fig. 3d**). In contrast, we found that _C_Ist2[590–936] did not associate with either type of liposome. Additional data showed that _C_Ist2[590–946] could substantially associate with liposomes containing only 5% PI(4,5)P_2_ but much more if these liposomes also included PS (**Fig. 3e**). We concluded that the Ist2 IDR efficiently binds to a membrane containing both PS and PI(4,5)P_2_ via its C-terminal polybasic motif, in support of previous results (23). Then, we mixed Osh6 with these PM-like liposomes in the absence or presence of _C_Ist2[590–946]. We observed that Osh6 associated significantly more with PM-like liposomes in the presence of the Ist2 IDR **(Fig. 3f)**. Collectively, these data confirm that Ist2 can associate with a membrane mimicking the inner leaflet of the yeast PM via its C-terminal end and further demonstrate that it can recruit Osh6 in this configuration.

### Osh6 localizes at artificial ER-PM contact sites scaffolded by the Ist2 IDR

Having established that _C_Ist2[590–946] can be attached to MPB-PE-containing liposomes via its N-terminal cysteine residue and bind to PS and PI(4,5)P_2_-rich liposomes via its C-terminal basic motif, we next sought to determine whether it could connect these two types of liposomes. For this purpose, we measured by DLS whether _C_Ist2[590–946] could induce the aggregation of L_A_ liposomes composed of DOPC and MPB-PE (90:10, 50 µM lipids) mixed with an equivalent amount of L_B_ liposomes composed of POPC, POPS and PI(4,5)P_2_ (65:30:5). Once mixed, in the absence of protein, the average hydrodynamic radius (R_H_) of these liposomes was approximately 100 nm with a low polydispersity (P_D_ < 50 nm). The addition of _C_Ist2[590–946] (500 nM) induced a substantial increase in the mean radius, reflecting its ability to promote liposome aggregation by connecting them (**Fig. 4a**). A more detailed analysis of the DLS measurements confirmed that, at the end of the kinetics, massive liposome aggregates form at the expense of free liposomes, with the appearance of a population with a mean radius ∼ 1000 nm (**Fig. 4b**). Control experiments with either L_A_ liposomes devoid of MPB-PE or L_B_ liposomes without PI(4,5)P_2_ showed, as expected, that no substantial aggregation occurred (**Figs. 4a** and **4b**). Next, we examined whether Osh6 impacted liposome tethering by Ist2 IDR. To facilitate this measurement, we used a construct (ATTO590-Osh6) devoid of free cysteine, meaning that no DTT had to be added to prevent the anchoring of Osh6 to the L_A_ liposomes. When ATTO590-Osh6 was added to L_A_ and L_B_ liposomes along with cIst2[590–946], we observed liposome aggregation to a level similar to that observed in our assays performed in the absence of Osh6 (**Fig. 4c**). We conclude that the Ist2 IDR can connect ER and PM-like membranes irrespective of the presence or absence of Osh6.

**Figure 4.**
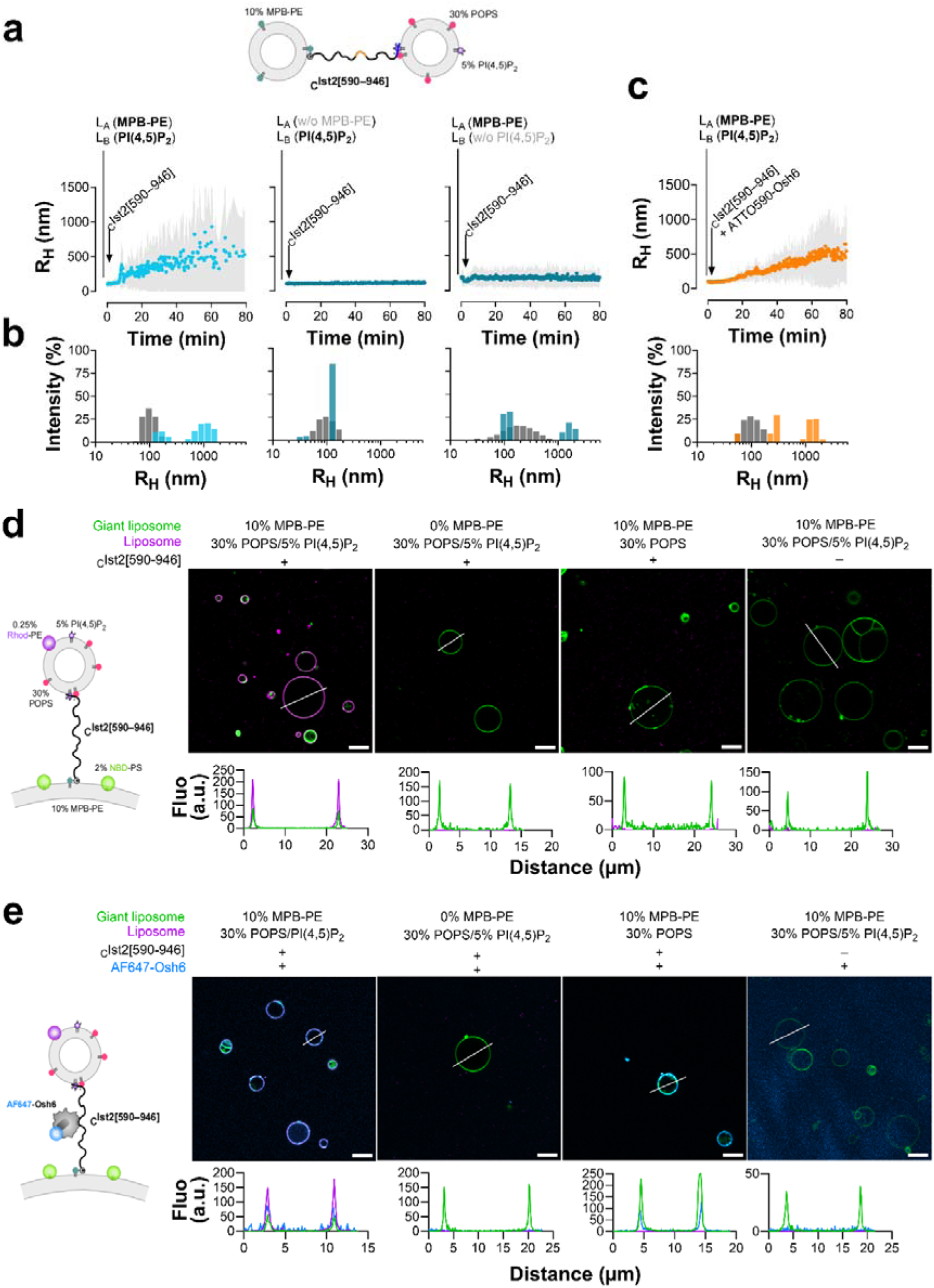
Ist2 can bridge ER- and PM-mimetic membranes and recruit Osh6. **(a)** DLS experiments. L_A_ liposomes composed of DOPC/MPB-PE/NBD-PE (90:10:0.1, 50 µM lipids) were mixed with an equivalent amount of L_B_ liposomes composed of POPC/POPS/PI(4,5)P_2_/NBD-PE (65:30:5:0.1) in HK buffer at 25 °C. Then, _C_Ist2[590–946] (0.5 µM) was mixed with liposomes. Control experiments were performed with L_A_ liposomes devoid of MPB-PE or L_B_ liposomes devoid of PI(4,5)P_2_. The mean radius (blue dots) and polydispersity (shaded area) of the liposome suspension were measured for 80 min. **(b)** Size distribution before (gray bars) and after the reaction (blue bars). **(c)** Aggregation kinetics measured with L_A_ liposomes containing MPB-PE and L_B_ liposomes containing PI(4,5)P_2_ upon addition of _C_Ist2[590–946] and ATTO590-Osh6 (0.5 µM). The mean radius (orange dots) and polydispersity (shaded area) of the liposome suspension were measured for 80 min. **(d)** Small liposomes with a PM-like composition are recruited at the surface of giant liposomes by the Ist2 IDR. Giant liposomes (∼65 µM accessible lipids) made of DOPC, doped with 2% NBD-PS, and containing or not 10% MPB-PE were incubated with small liposomes (166 µM accessible lipid), composed of 70% POPC, 30% POPS and 0.25% Rhod-PE, doped or not with 5% PI(4,5)P_2_ (at the expense of POPC), in the absence or the presence of 1 µM _C_Ist2[590–946]. All images were obtained by confocal microscopy (Leica TCS SP8, 63×, NA 1.4). The line scans show fluorescence intensities of the green and magenta channels along the white line. Scale bar, 10 µm. **(e)** Osh6 is recruited at the level of PM-like and ER-like membranes connected by the Ist2 IDR. AF647-Osh6 (80 nM) was added to giant liposomes composed of DOPC/MPB-PE/NBD-PS (88:10:2), mixed with small liposomes composed of POPC/POPS/Rhod-PE (70:30:0.25), doped or not with 5% PI(4,5)P_2,_ and _C_Ist2[590–946]. A control experiment was carried out without _C_Ist2[590–946]. The line scans show fluorescence intensities of the green, magenta, and blue channels along the white line. Scale bar, 10 µm.

To further validate our approach for reconstituting artificial ER-PM contacts with Osh6:Ist2 complexes, we visually examined whether Osh6 could be at the interface between ER- and PM-like membranes connected by the Ist2 IDR. We prepared giant liposomes composed of DOPC/MPB-PE/NBD-PS (88:10:2, ∼60 µM lipids) and incubated them with an equivalent amount of small liposomes composed of POPC/POPS/PI(4,5)P_2_/Rhod-PE (65:30:5:0.25), in the presence of _C_Ist2[590–946] (1.5 µM). Observation by confocal microscopy revealed that the surface of the green giant liposomes was systematically coated with small red liposomes, as analyzed by line-scan analyses (**Fig. 4d**). The absence of either MPB-PE in the membrane of giant liposomes, PI(4,5)P_2_ in the membrane of small liposomes or _C_Ist2[590–946] resulted in no recruitment of small liposomes to giant liposomes. Therefore, these data indicate that small liposomes with a PM-like composition can be recruited to the surface of giant liposomes coated with the Ist2 IDR. Remarkably, upon addition of AF647-Osh6 to giant liposomes functionalized with _C_Ist2[590–946] and mixed with PI(4,5)P_2_-containing small liposomes, we observed far-red fluorescence overlapping with red and green fluorescence (**Fig. 4e**). The absence of MPB-PE in the membrane of giant liposomes or that of _C_Ist2[590–946] disrupted Osh6 recruitment. We concluded that Osh6 can localize between ER- and PM-like membranes connected by the Ist2 IDR.

### Ist2 IDR ensures a fast and directed Osh6-mediated PS transfer between connected membranes

The successful reconstitution of Osh6 recruitment to artificial ER-PM contact sites prompted us to investigate the ability of Osh6 to transfer PS from ER- to PM-like membranes using a fluorescence assay based on NBD-PS. As the composition of our PM-like liposomes included POPS and PI(4,5)P_2_, we anticipated that these lipids, as natural and fortuitous ligands, respectively, might interfere with the NBD-PS transfer activity of Osh6 based on our previous work (44). To mitigate this, we resorted to the Osh6(HH/AA) variant, less prone than the wild-type Osh6 to transfer PI(4,5)P_2_ (**Fig. S5a**). GST pull-down and flotation assays showed that this construct binds well to the Ist2 IDR (**Fig. S5b-d**). To perform the NBD-PS transfer assay, we mixed L_A_ liposomes containing MPB-PE and NBD-PS with L_B_ liposomes enriched with PS and doped with 2% Rhod-PE and 5% PI(4,5)P_2_, and L_C_ liposomes only enriched with PS. Next, the _C_Ist2[590–946] construct was added to the sample to connect the L_A_ with the L_B_ liposomes. Upon adding Osh6(HH/AA), we measured a fast transfer of NBD-PS from L_A_ to L_B_ liposomes in the presence of L_C_ liposomes, as indicated by an increase in FRET between NBD-PS and Rhod-PE (**Fig. 5a, trace i**). The same experiment, but with Rhod-PE in L_C_ instead of L_B_ liposomes, showed that NBD-PS transfer from L_A_ to L_C_ liposomes was much slower than to L_B_ liposomes (**Fig. 5a, trace ii**). The prominent role of the Ist2 IDR in the Osh6-mediated lipid transfer was further substantiated by the fact that in its absence, NBD-PS was evenly transferred to L_B_ and L_C_ liposomes (**Fig. 5a, traces iii and iv**). When using Osh6 WT instead of Osh6(HH/AA), we measured lower NBD-PS transfer rates in all conditions, in line with the idea that PI(4,5)P_2_ might be trapped by Osh6 and interfere with its NBD-PS transfer activity, but we observed the same trend (**Fig. S5e,f**). On the contrary, with the ORD of ORP8 (bearing an HH/AA double mutation), we did not observe any acceleration or preference of transport toward L_B_ liposomes connected to L_A_ liposomes by _C_Ist2[590–946] (**Fig. 5b, c**). These results suggest that the specific association of Osh6 with the disordered region of Ist2 allows for a rapid and accurate PS transfer between membranes connected by Ist2, without mistargeting toward a third compartment.

**Figure 5.**
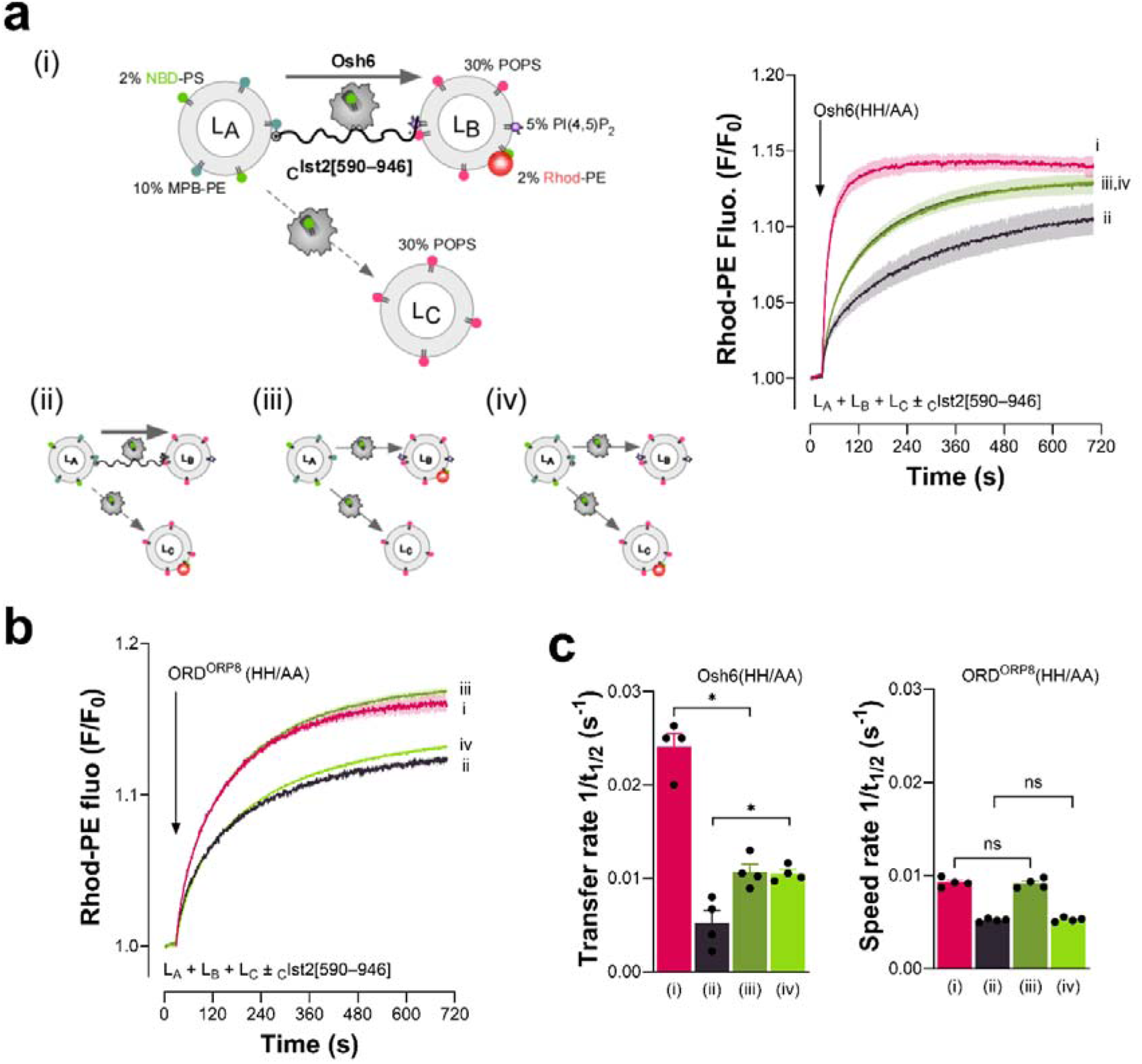
Tethering of membranes by Ist2 IDR promotes fast and directed flux of PS by Osh6. **(a)** Real-time PS transport assays. (i) Osh6(HH/AA) was added to L_A_ liposomes (200 µM), doped with MPB-PE and 2% NBD-PS, and connected by cIst2[590–946] (0.5 µM) to L_B_ liposomes (200 µM), containing PS, PI(4,5)P_2_ and 2% Rhod-PE in the presence of free L_C_ liposomes (200 µM) with no PI(4,5)P_2_. The fast increase in rhodamine fluorescence (λ_ex_ = 460 nm, λ_em_= 580 nm) over time corresponds to NBD-PS transfer from L_A_ to L_B_ liposomes. A mirror experiment (ii), in which Rhod-PE is incorporated in L_C_ and not L_B_ liposomes, was conducted to measure the specific transfer of PS from L_A_ to L_C_ liposomes. In the absence of _C_Ist2[590–947], Osh6(HH/AA) transfers equally PS to L_B_ and L_C_ liposomes (iii, iv). Each curve represents the mean ± s.e.m. of several kinetics recorded during independent experiments (n = 3). **(b)** ORD^ORP8^(HH/AA) does not preferentially transfer PS between Ist2 IDR-connected membranes, as there is no association between ORD^ORP8^ and Ist2 IDR. **(c)** Rate (1/*t*_1/2_) of NBD-PS transfer measured with Osh6(HH/AA) and ORD^ORP8^(HH/AA) in conditions (i), (ii), (iii), and (iv).

### Osh6 mutations that prevent interaction with the Ist2 IDR hinder Osh6-mediated PS transfer

To further dissect the role of the Ist2 IDR in Osh6-mediated PS transfer, we designed Osh6 mutants that could not bind to it. To this end, we used AlphaFold3 (49) to build a model of the Osh6:Ist2 IDR complex, using *S. cerevisiae* Osh6 and Ist2[590–946] sequences. Remarkably, we obtained five top-ranked models (ranking score 0.98–0.99) where the [719–750] segment of Ist2, encompassing the minimal Osh6-binding motif, was in close contact with Osh6’s surface **(Fig. 6a, Fig. S6a)**. Moreover, based on the Predicted Aligned Error (PAE, **Fig. S6b**), the values predicted by the local distance difference test (pLDTT) assigned to each Ist2 residues (**Fig. 6a, Fig. S6a)**, and the ipTM score (0.79 for each model, **Fig. S6a**), we concluded that the position and orientation of the [720–770] segment of Ist2 IDR relative to Osh6, the folding of the Osh6-binding motif, and the Osh6:Ist2 binding interface were predicted with high confidence. (49). An in-depth analysis of these models suggested that the [719–750] segment of Ist2 associates with an Osh6 surface region composed of residues belonging to the β3-β4 and β5-β6 loops, and two helices, α8 and α9, which form the C-terminal end of the protein. Of note, this region lies opposite the entrance of the lipid-binding pocket of Osh6. Interestingly, in all models, we noted that two Osh6 residues (Q417 and Q418) are engaged in multiple H-bonds with the T736 residue in Ist2, previously found to be critical for the Osh6:Ist2 interaction (16) **(Fig. 6b, c, Sup. Excel Table)**. Moreover, for all five models, we observed that the backbone and/or side chains of other Osh6 residues (E170, Y391, Q409, T420, F419, K722) are engaged in H-bonds with the same specific residues within the Ist2 IDR. Remarkably, in three models, the side-chain of the D141 residue in Osh6, found to be necessary for Osh6:Ist2 interaction (16), is engaged in an H-bond with the side-chain of the R750 residue in Ist2 (**Sup. Excel Table)**. Thus, we obtained a model of Osh6:Ist2 IDR complex in good agreement with previous binding data (16, 17) and identified Osh6 residues, in addition to D141 (16), that might be key for interacting with Ist2.

**Figure 6.**
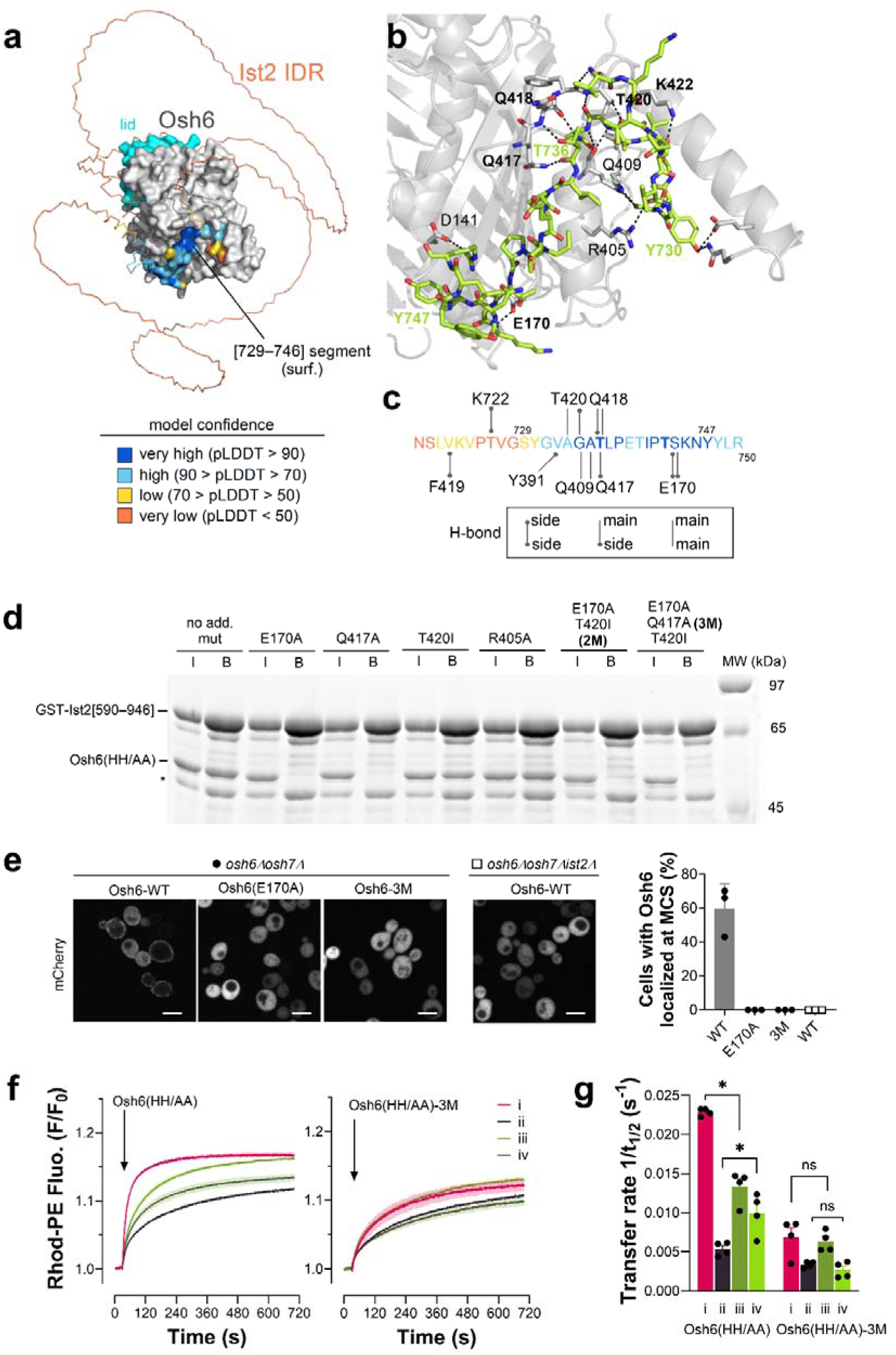
Identification of Osh6 mutants unable to preferentially transfer PS between Ist2-connected membranes. **(a)** AlphaFold3 model of the Osh6:Ist2 IDR complex. Osh6 is represented as a grey surface. The molecular lid, gating the lipid-binding pocket of the protein, is colored in cyan. For clarity, the N-terminal low complexity sequence of Osh6 (35 residues) has been omitted. Ist2 IDR is represented in ribbon mode with the Ist2[729–747] segment (minimal Osh6-binding motif) represented as a surface. Ist2 IDR is colored according to the predicted local distance difference test (pLDDT) *per* atom. **(b)** Close-up view of the potential H-bonds network (dark dashed lines) between Osh6 and the Ist2 IDR. The Ist2[719–750] segment is represented in stick mode with carbons in green, nitrogens in blue, and oxygens in red. Reference residues are labeled in green. Osh6 is represented in transparent cartoon mode. The side and/or main chain of Osh6’s residues involved in H-bonds with Ist2’s residues are represented in stick mode (carbons in grey) and labeled in black (in bold: residues systematically involved in H-bonds with Ist2 residues in all five AlphaFold3-predicted models; D141 and R405 are example of Osh6’s residues that are not systematically engaged in H-bonds with Ist2 residues from one model to the other). **(c)** H-bonds network between Osh6 and the Ist2 IDR. The sequence of the [719–750] segment of Ist2 encompassing the minimal Osh6-binding motif (residue 729 to 747) is colored according to the pLDDT. Osh6’s residues’ names and positions are in black. The H-bonds between the main chain and side chains of Ist2’s and Osh6’s, systematically observed in the five AlphaFold3-derived models, are represented as grey lines. **(d)** GST pull-down with Osh6 mutants. GST or GST-Ist2[590–946] was immobilized on beads and incubated with Osh6(HH/AA) construct with or without additional mutations (E170A, Q417A, T420I, R405A, E170A/T420I (2M), E170A/Q417A/T420I (3M)). Input (I) and bound (B) fractions were analyzed using SDS-PAGE. **(e)** Localization of Osh6 variants (Osh6-WT, Osh6[E170A] and Osh6[E170A/Q417A/T420I] (3M)), expressed from a low-copy (CEN) plasmid and tagged C-terminally with mCherry, in *osh6*Δ*osh7*Δ cells. Localization of Osh6-WT in *osh6*Δ*osh7*Δ*ist2Δ* cells is shown for comparison. Scale bar, 5 µm. The quantification of the percentage of cells with Osh6 localized at contact sites in *osh6*Δ*osh7*Δ cells is shown on the right (mean of n ≥ 100 cells, analyzed in 3 independent experiments). **(f)** Real-time PS transport assays. (i) Osh6(HH/AA) or Osh6(HH/AA)-3M (200 nM) was added to L_A_ liposomes doped with MPB-PE and NBD-PS and connected by cIst2[590–946] (0.5 µM) to L_B_ liposomes containing PS, PI(4,5)P_2_ and Rhod-PE in the presence of free L_C_ liposomes devoid of PI(4,5)P_2_. The increase in fluorescence over time corresponds to NBD-PS transfer from L_A_ to L_B_ liposomes. A mirror experiment (ii) was conducted to measure the specific transfer of NBD-PS from L_A_ to L_C_ liposomes. These two experiments were also performed without _C_Ist2[590–946] (iii, iv). Each curve represents the mean ± s.e.m. of several kinetics recorded during independent experiments (n = 4). **(g)** Rate (1/*t*_1/2_) of NBD-PS transfer measured with Osh6(HH/AA) and mutants in conditions (i), (ii), (iii), and (iv).

Based on AlphaFold3 predictions, we decided to mutate individually or in a combined manner Osh6 residues (E170, Q417, T420) that, in all the models, form via their side-chain H-bonds with Ist2 residues (G734, T736, S743) whose position, at the Ist2:Osh6 interface, was predicted with the highest confidence (pLDDT value> 90, **Fig. 6c**). We also selected a residue (R405) whose side-chain was predicted to be involved in an H-bond with Ist2 in only one model (**Sup. Excel Table)**. Therefore, we purified Osh6(HH/AA) constructs with one additional mutation (E170A, or Q417A, or T420I, or R405A), double (E170A/T420I called 2M) or triple mutations (E170A/Q417A/T420I, called 3M, **Fig. S6c**). All these mutants were properly folded, as judged by their CD spectra (**Fig. S6d**). In line with our predictions, we found that Osh6(HH/AA) constructs bearing an E170A or Q417A mutation, but also the Osh6(HH/AA)-2M or Osh6(HH/AA)-3M construct, were unable to bind to the Ist2 IDR (**Fig. 6d**). Conversely, the mutation of T420 or R405 did not impact the association of Osh6 with Ist2. To confirm these results at the cellular level, we analyzed the localization of Osh6(E170A) and Osh6-3M variants, chromosomally tagged with mCherry, and found that they were fully cytosolic. In contrast, Osh6 WT was mostly observed in patches at the cell cortex, indicative of its presence at ER-PM contact sites (15–17, 45) **(Fig. 6e).** Control experiments showed that deletion of Ist2 rendered Osh6 WT cytosolic, as observed for the mutants when Ist2 is present. We conclude that residues E170 and Q417 are critical for Osh6 to recognize the Ist2 IDR. Finally, we examined the ability of the 3M mutant to preferentially transfer NBD-PS between L_A_ and L_B_ liposomes connected by _C_Ist2[590–946] in the presence of free L_C_ liposomes. We found that contrary to Osh6(HH/AA), Osh6-3M did not preferentially transfer PS between liposomes connected by the Ist2 IDR; in all conditions tested (with or without _C_Ist2[590–747]), its transfer activity was similar to that of Osh6(HH/AA) measured between unconnected liposomes (**Fig. 6f, g**). Taken together, these data not only validate our structural predictions of the Osh6:Ist2 interface but also confirm the specific requirement of the Osh6:Ist2 interaction for a fast and accurate, directed PS transfer between the ER and the PM.

### Osh6-mediated PS transfer can be coupled with the PS-scrambling activity of Ist2

We have recently shown that Ist2 is a lipid scramblase at the ER (28). Therefore, a potential scenario is that Ist2, by equilibrating PS (among other phospholipids) between the leaflets of the ER membrane, fuels Osh6 for the transfer of PS to the PM. However, we did not observe any impact of the deletion of the Ist2 transmembrane scrambling domain on the cellular PS levels or distribution (28). One issue may be the sensitivity/specificity of our assays. Indeed, since deletion of the Ist2 transmembrane domain is synthetically lethal with Cho1 deletion, we cannot implement our PS transfer assay in the ΔNIst2 strain. To explore whether Ist2 and Osh6 can partner for the redistribution of lipids between the ER and the PM, we measured the transfer of NBD-PS by Osh6 from proteoliposomes (PLs), reconstituted with the N-terminal part of Ist2, to liposomes mimicking the PM. First, we purified full-length Ist2 (hereafter called Ist2) and Ist2[1–600] (Ist2ΔC) fused to a biotin-acceptor domain after overexpression in *S. cerevisiae*. The two constructs were then reconstituted into PLs made of DOPC and doped with 2 mol% of NBD-PS, i.e., with an ER-like composition identical to the PS donor liposomes we used so far in our transfer assays. For all conditions, most of the purified Ist2 was recovered after detergent removal with Bio-beads **(Fig. S7a).** DLS indicated that the average R_H_ was ∼61, 56, and 64 nm for Ist2, Ist2ΔC, and mock PLs, respectively **(Fig. S7b)**. Then, we conducted scramblase assays in which dithionite was used to selectively extinguish the fluorescence of NBD-labelled lipid present in the outer leaflet of the PLs (50). Adding dithionite to “mock” PLs provoked a ∼63% reduction in the NBD signal, meaning that the same percentage of NBD-PS was accessible to the reducing agent. With Ist2 and Ist2ΔC PLs, the fluorescence was reduced by 81 and 76%, respectively, indicating that more NBD-PS was accessible on the outer leaflet **(Fig. 7a, c)**. These values, although lower than those obtained when Ist2 and Ist2ΔC were reconstituted in POPC/POPS PLs **(Fig. S7a,c)**, as in our previous study (28), indicated that Ist2 and Ist2ΔC exhibit a robust scramblase activity in ER-like membranes.

**Figure 7.**
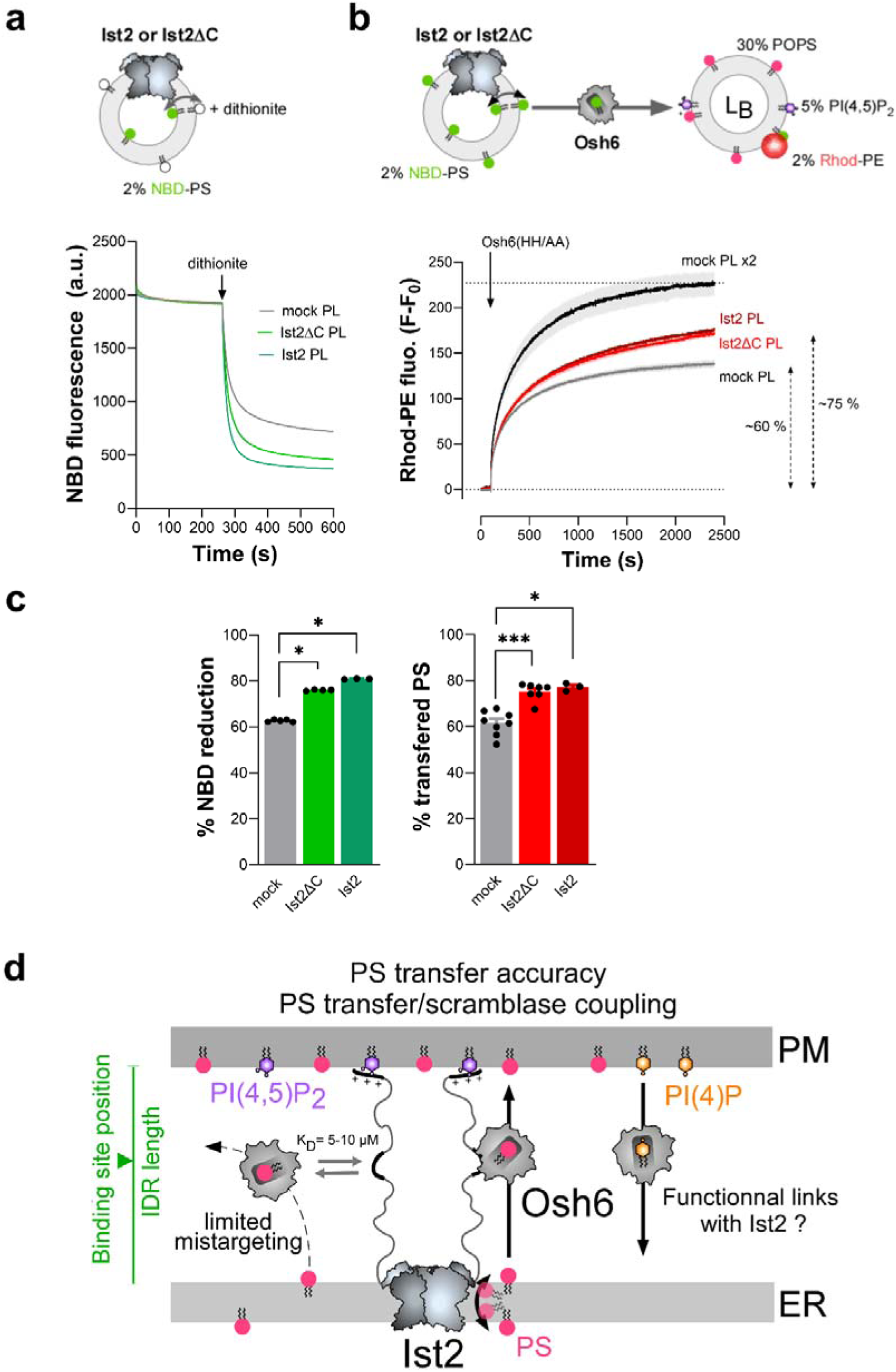
Coupling between the PS scrambling activity of Ist2 and the PS transfer activity of Osh6. **(a)** Scramblase assay. NBD-PS-containing proteoliposomes (PLs), in which the Ist2ΔC or Ist2 construct is reconstituted, or protein-free liposomes (“mock”), have been diluted in HK buffer at 25 °C under constant stirring. The amount of NBD-PS is the same in the three conditions, as indicated by the NBD fluorescence level. After 5 min, 10 mM of dithionite was injected, and the quenching of NBD was measured over an additional 5 min. Each curve represents several kinetics’ mean ± s.e.m. (n = 4). **(b)** Real-time PS transport assays. Proteoliposomes, composed of DOPC/NBD-PS (98:2), and reconstituted with the Ist2ΔC or Ist2 construct, or protein-free liposomes (“mock”), were mixed with 50 µM of L_B_ liposomes composed of POPC/POPS/PI(4,5)P_2_/Rhod-PE (63:30:5:2) in HK buffer at 25 °C. An equal amount of NBD-PS was present in the three conditions. Then, Osh6(HH/AA) was added, and NBD-PS transport from proteoliposomes to L_B_ liposomes was followed over time by measuring FRET at 580 nm (excitation λ = 460 nm). PS transfer is expressed as F–F_0_, F_0_ being the fluorescence measured before adding Osh6. The final signal for each condition was then normalized to the final signal (F_max_) measured in PS transfer assays, where twice the amount of mock liposomes was used to mimic a condition where 100% of PS would be accessible at the surface of a single amount of mock, Ist2, or Ist2ΔC PLs. Each curve represents the mean ± s.e.m. of several kinetics (n = 3–8). **(c)** Percentage of NBD reduction by dithionite and relative percentage of Osh6-mediated PS transfer, derived from the data shown in (a) and (b) as a function of the type of proteoliposome used in each assay. Unpaired Mann–Whitney U test (\**p* < 0.05, \*\*\**p* < 0.001). **(d)** Model of the functional partnership between Osh6 and Ist2. Osh6 can bind to the Ist2 IDR in a PS-bound state and thereby transfer PS from the ER to the PM in a fast and directed manner, with a limited capacity to deliver PS by mistake into another cellular membrane. Moreover, the PS transfer activity of Osh6 can be coupled with the PS scrambling activity of Ist2. The length of the Ist2 IDR and the position of the Osh6-binding site in it are critical factors that impact how efficient Osh6-mediated PS transfer is. Whether the association of Osh6 with Ist2 has a functional impact on its ability to transfer PI(4)P from the PM to the ER has to be investigated.

Having established the activity of Ist2, we measured the PS transfer assay of Osh6 added to “mock”, Ist2ΔC or Ist2 PLs (∼50 µM lipids) pre-incubated with a similar amount of L_B_ liposomes composed of POPC/POPS/PI(4,5)P_2_/Rhod-PE (63:30:5:2). Importantly, the amount of NBD-PS was the same in all conditions (**Fig. S7d).** With “mock” PLs, we measured a rapid increase in FRET from NBD-PS to Rhod-PE, reflecting the transfer of NBD-PS from PLs to L_B_ liposomes, which reached near equilibration after 1 h **(Fig. 7b)**. Strikingly, when Osh6 was added to Ist2ΔC or Ist2 PLs, we measured a higher increase in FRET signal within the first minutes; moreover, this signal kept increasing even after 1 hour without reaching a plateau. These kinetics data suggested that the scramblase activity of Ist2 and Ist2ΔC sustained NBD-PS transfer by Osh6 to L_B_ liposomes.

To strengthen these results, we first assessed by a collisional quenching assay that ∼50% of the NBD-PS in the membrane of mock, Ist2ΔC, or Ist2 PLs was protected from iodide, indicating that this lipid was distributed equally between the membrane leaflets of PLs for each preparation (**Fig. S7e**). Second, we verified that Osh6 did not affect the scramblase activity of Ist2 **(Fig. S7f)**. Third, we mimicked a situation where all the PS molecules would already be in the outer leaflet of the PLs before adding Osh6. To do so, we measured PS transfer using twice the amount of “mock” PLs and observed a transfer of PS higher than that observed with FL-Ist2 and Ist2ΔC PLs within a few seconds **(Fig. 7b)**. This meant that the effect seen with Ist2 and Is2ΔC PLs was not due to a pre-existing availability of PS that would be higher with these PLs but to the scramblase’s capacity to replenish the PS pool in the outer leaflet of PLs while Osh6 functioned. By normalizing the PS transfer data, we found that Osh6 transferred up to ∼15% more PS from Ist2/Ist2ΔC PLs than from “mock” PL, which was remarkably in line with the dithionite assay showing that up to ∼20% more NBD-PS was accessible due to the Ist2 scrambling activity **(Fig. 7c)**. From these experiments, we conclude that Ist2 can augment the PS pool accessible to Osh6 in ER-like membranes and suggest that the PS transfer activity of Osh6 may be coupled to the Ist2 scramblase activity.

## Discussion

An important and emerging issue in cell biology is to understand why LTPs are localized to contact sites and how their concerted action with other lipid transporters and membrane tethering factors affects intracellular lipid distribution and metabolism. Here, we dissect *in vitro* how Osh6 associates with the IDR of Ist2. Using cell-based assays, we also report that the length of the Ist2 IDR and the position of the Osh6 binding site are essential for PS transport at ER-PM contact sites. We recapitulate PS transfer from the ER to the PM using purified components and determine that the association of Osh6 with Ist2 dramatically increases the efficiency of PS transfer between membranes connected by the Ist2 IDR. Moreover, we demonstrate a functional partnership between the PS transfer activity of Osh6 and the scramblase activity of Ist2. These results allow us to propose a model on the functional partnership between Osh6 and Ist2 (**Fig. 7d**).

First, we quantified how Osh6 associates with a purified form of the full-length Ist2 IDR, which we confirmed has no secondary structure and a hydrodynamic radius close to that predicted or obtained for an IDR of similar length (e.g., the 351 aa-long IDR of GHR-ICD with an R_H_ = 5.08 nm) in solution (51, 52). The affinities of Osh6 for the full-length IDR or a peptide encompassing the Osh6 binding site were similar (K_D_ = 6–9 µM), and close to those measured by Dutzler and co-workers between Osh6 and short segments of Ist2 IDR, using calorimetry (K_D_ = 0.71–1.1 µM) (29). This suggests that Osh6 has no difficulty reaching its binding site at the center of the Ist2 IDR, i.e., in the core of a region that likely forms a molten globule, as suggested by our structural models. Moreover, regardless of whether it is empty or loaded with a lipid ligand, Osh6 interacts with the Ist2 IDR similarly, and this interaction does not impact how Osh6 traps lipid ligands. This means that Osh6 can operate lipid exchange cycles while interacting with Ist2 and keeping its localization at contact sites.

What can the measured affinity of Osh6 for Ist2 tell us about the association of the two proteins in the yeast cell? The median volume of a budding yeast cell is 44 fL, corresponding to a sphere of 4.4 µm in diameter (53). For a yeast of this size, the PM area is estimated to measure 102 µm^2^ (54). Considering 4,100 Ist2 copies/cell (https://www.yeastgenome.org/) and all the Ist2 proteins are at ER-PM contact sites, i.e., between two membranes that are 20 nm apart (19), the local Ist2 concentration there could be about 3.4 µM. In contrast, the cytosolic concentration of Osh6 (3,600 copies/cell) is 135 nM. Considering a 1:1 binding model and a K_D_ between 1–10 µM, the fraction of Ist2-bound Osh6 in ER-PM contact sites could be 25–75% at a steady state. This fits well with the observation that Osh6 is partly cytosolic and at the cortical ER (15–17, 45). Importantly, by modifying the length of the Ist2 IDR or the position of the Osh6-binding site in the Ist2 IDR, we conclude that a minimal length of the IDR and/or distance of the Osh6-binding site from the ER is sufficient to support PS transport. Of note, we show that the attachment of the IDR to two membranes is required for its proper localization, because deletion of either the TM domain or the PM-binding region of Ist2 leads to its aggregation in cells, precluding further analysis. However, more subtle defects may be missed by these assays due to limitations in their sensitivity. For example, it was shown that IDRs present in the mammalian Osh-related LTPs ORP4 and OSBP influence their mobility within contact sites (55). Nevertheless, our results suggest that the affinity and thus association time between Osh6 and the Ist2 tail exceeds the time necessary for an intermembrane lipid transfer event. Otherwise stated, thanks to its affinity for Ist2, Osh6 might function as an LTP capable of simultaneously transferring lipid and tethering membranes (e.g., ORP5/8) at contact sites. Additional mechanistic investigations are needed to address this point.

Further, we reconstituted ER-PM contact sites by anchoring the Ist2 IDR via its N-terminus to neutrally charged liposomes with low lipid packing, mimicking the ER membrane, and by adding them to liposomes with higher lipid packing, rich in PS with a minute amount of PI(4,5)P_2_, mimicking the PM. In line with previous reports (23, 24), we observed that the membrane-binding capacity of the C-terminal motif of the Ist2 IDR strongly depends on PI(4,5)P_2_, which explains why Ist2 specifically anchors the ER to the PM. Next, we established that Osh6 could be recruited to these artificial contact sites. Interestingly, in the absence of the Ist2 IDR, Osh6 binds weakly to ER and PM-like membranes. This suggests that, even when associated with Ist2, Osh6 would maintain its ability to associate transiently with these membranes, which is necessary for its transfer activity (45).

Once able to reconstitute Osh6:Ist2 complex at contact sites and knowing the strong selectivity of Ist2 IDR for PI(4,5)P_2_, we could then measure Osh6-mediated PS fluxes between ER and PM-like liposomes (containing PS and PI(4,5)P_2_) connected by Ist2 in the presence of a third population of liposomes, devoid of PI(4,5)P_2_. In this experimental setup that recapitulates the multi-compartmentalized organization of a cell, we found that the specific association of Osh6 with the Ist2 IDR allows for a fast transfer of PS between connected membranes while preventing Osh6 from transferring PS toward another compartment. In the absence of interaction, PS was slowly transferred to all membranes. This matches well our observation that the lack of Ist2 or Ist2:Osh6 association leads to a defect in the delivery of PS by Osh6 into the PM (16); likely, PS is slowly spread by Osh6 toward different organelles and membranes. Together, our data show that the localization of LTPs at contact sites allows for a rapid and accurate lipid transfer between two membranes in the complex environment of the cell.

So far, in vitro studies focused on LTPs (OSBP, STARD3, ATG2A) with a dual ability to tether membranes and carry lipids, showing that they efficiently transfer lipids between liposomes once they connect them (56–59). In other studies, lipid-transfer modules anchored to donor liposomes delivered lipids efficiently to acceptor liposomes only when both liposomes were brought nearby by a tethering factor (60–62). Consequently, it was impossible to distinguish whether lipid transfer was enhanced due to the concentration of LTPs between membranes, a shorter intermembrane distance reducing the time needed for a LTP’s transfer module to move lipids from one membrane to the other, or just because the lipid-transfer module had access to the two membranes. By contrast, our system is unique as Osh6 is neither attached to the membrane nor has a membrane tethering capacity, which is a function ensured by Ist2. We could thus demonstrate that mutations of Osh6 that impair its binding to the Ist2 IDR (Osh6-3M) reduce the PS transfer rate in otherwise intact synthetic ER-PM contact sites, implying that a closer proximity between membranes is not sufficient to confer a kinetic advantage. This aligns with our observations that Osh6-mediated PS transfer is strongly impaired in yeast expressing a mutant form of Ist2, unable to recruit Osh6 but still able to form ER-PM contact sites (16). Thus, our data suggest that LTPs can work optimally at the contact sites because they are concentrated in these substructures and not because they benefit from a short intermembrane space.

Our studies also provide structural insights into how Osh6 binds to Ist2 by experimentally validating models of the Osh6:Ist2 complex generated by AlphaFold3 (49). Remarkably, the algorithm managed to identify the 727–750 segment in the full-length Ist2 IDR as the Osh6 binding motif and provided models in line with our previous results obtained by two-hybrid assays, pointing to the contribution of Osh6’s residues (D141 and L142) and Ist2’s residues (T736) to Osh6:Ist2 interaction (16). Moreover, these models allow to identify several Osh6’s residues (E170, Y391, Q417, T420) that are engaged in H-bonds with Ist2 residues, as also indicated by the structures of Osh6 recently solved in complex with Ist2[732–761] and Ist2[732–757] peptides (29). We confirmed the relevance of these models by testing Osh6 mutants, showing that Q417 and E170 residues are critical for the Osh6:Ist2 interaction. Importantly, our data provide evidence that the Ist2 binding site on the Osh6 surface is distal to the entrance of the lipid-binding pocket, thereby explaining why Osh6 can transfer lipids when it binds to the Ist2 IDR. Interestingly, the β-barrel that constitutes the core of the ORD, hosting lipid ligands, is a highly conserved structure among ORP/Osh proteins, but it is decorated, notably at its C-terminus, by structural elements that strongly differ depending on ORP/Osh subfamilies (6). The fact that Osh6 associates with Ist2 mainly via its C-terminal region establishes that these structural elements allow for a specific interaction between an ORD and a partner. Whether this is true for other ORP/Osh proteins remains to be defined.

Recently, we and others have unveiled that the transmembrane domain of Ist2 has a scramblase activity (28, 29). Thus, in addition to its role as ER-PM tether and Osh6 binding partner, Ist2 equilibrates lipids between the leaflets of the ER membrane, suggesting a possible coupling between a transbilayer and intermembrane transport mechanism. We tested this idea in vitro, showing that Ist2, by scrambling PS between the leaflets of ER-like membranes, sustains the transfer of PS from these liposomes by Osh6 to PM-like membranes, likely by constantly replenishing the PS pool at the surface of ER-like membranes. Therefore, we established that the coordinated work of a scramblase and an LTP can ensure lipid flux between a membrane’s inner leaflet and another membrane’s outer leaflet. However, our first data suggest that the Ist2 scrambling activity does not promote Osh6-mediated PS transfer activity in yeast (28). Thus, the coupling between Ist2 and Osh6 observed in vitro might serve another function in vivo.

Interestingly, it is assumed that scramblases could prevent the membranes from being destabilized by the delivery or extraction of lipids by LTPs, via the re-equilibration of lipids between the membrane leaflets. Such a mechanism of compensation has been proposed for autophagosome formation, as LTP ATG2 can associate with the ER-resident scramblases TMEM41B/VMP1 but also the ATG9 scramblase present in the nascent autophagosome (38). Likewise, the LTP VPS13A and the scramblase XK, whose mutations cause pathological conditions with similar manifestations and both of which are essential for PS exposure in ATP-stimulated cells, have been shown to interact at ER-PM contacts (63, 64). Therefore, one scenario is that Ist2, by transporting PS but also PC and PE (29) across the ER membrane, maintains the integrity and/or proper lipid composition of the two leaflets of this membrane when PS is delivered to the PM by Osh6. Besides, we still do not know to what extent the coupling between the transport activity of Osh6 and Ist2 might depend on their physical association. We did not observe differences in Osh6 transfer activity using Ist2 and Ist2ΔC proteoliposomes, but the concentration of Ist2 was much lower than that of Osh6 in our assays, which prevents conclusions. Thus, additional in vitro and in-cell investigations are needed to fully determine how exactly the lipid transport activities of Osh6 and Ist2 are coupled at ER-PM contact sites and impact cellular functioning.

## Material and Methods

### Protein expression, labeling, and purification from *E. coli*

Osh4, Osh6, Osh6(noC/S190C), Osh6(H157A/H158A), and other Osh6 mutants, ORD^ORP8^ and ORD^ORP8^ (H514A/H515A), NBD-PH_PLCδ1_ were purified as previously described (44, 65). To obtain Osh6 labeled with an ATTO590 maleimide (ATTO-TEC) or an Alexa Fluor 647-C2 maleimide dye (Invitrogen), an Osh6(noC/T262C) mutant was produced, labeled with the respective dye and purified following the protocol in (45). The concentration of all these constructs was determined by UV spectrophotometry.

GST-Ist2[590–936], GST-Ist2[590–946], GST-_C_Ist2[590–936], GST-_C_Ist2[590–946], GST-Ist2[590–946]Δ727–749, GST-Ist2[590–768], and GST-Ist2[727–776] were expressed in *E. coli* (BL21-GOLD(DE3)) competent cells (Stratagene) grown in Luria Bertani Broth (LB) medium at 30 °C overnight upon induction with 1 mM isopropyl β-D-1-thiogalactopyranoside (IPTG). When the optical density of the bacterial suspension, measured at 600 nm (OD_600_), reached a value of 0.6—0.7, bacterial cells were harvested and re-suspended in cold TN buffer (50 mM Tris-HCl, pH 7.4, 150 mM NaCl, 2 mM DTT) supplemented with 1 mM PMSF, 10 µM bestatin, 1.6 µM pepstatin A and cOmplete®, EDTA-free protease inhibitor tablets (Roche). Cells were lysed in a Cell Disruptor TS SERIES (Constant Systems Ltd.), and the lysate was centrifuged at 186,000 × *g* for 1 h 30. Then, the supernatant was applied to Glutathione Sepharose 4B (Cytiva) for 3 h 30 at 4 °C. To purify the Ist2[590–936], _C_Ist2[590–936], Ist2[590–946] and _C_Ist2[590–946] constructs, the beads were washed four times with TN buffer devoid of protease inhibitors; the beads were incubated with thrombin overnight at 4 °C to cleave off the Ist2 construct from the GST domain. Each construct was recovered in the supernatant after several cycles of centrifugation and washing of the beads, concentrated, and injected onto an XK-16/70 column packed with Sephacryl S-200 HR to be purified by size-exclusion chromatography. The fractions with ∼100% pure Ist2 construct were pooled, concentrated, and supplemented with 10% (v/v) pure glycerol (Sigma). Aliquots were prepared, flash-frozen in liquid nitrogen, and stored at -80 °C. The protein concentration was determined using a BCA assay. For some experiments, a volume of 100 µL from a stock solution of _C_Ist2[590–946] was applied onto a 0.5 mL Zeba spin desalting column (7 kDa molecular weight cut-off) equilibrated with freshly degassed HK buffer, according to manufacturer’s indications, to remove DTT from the protein, and immediately used. For some spectroscopy measurements, the concentration of the _C_Ist2 constructs was determined by measuring their absorbance at λ = 280 nm (ε = 81,820 M^-1^.cm^-1^). To purify the GST-Ist2[590–946], GST-Ist2[590–946]Δ727–749, GST-Ist2[590–768], and GST-Ist2[727–776] constructs, the beads were washed four times with TN buffer devoid of protease inhibitors; then the beads were incubated with TN buffer containing 20 mM glutathione and 2 mM DTT. The protein concentration was determined using a BCA assay.

ORD^Osh3^ (Osh3[605-996]) was produced in fusion with an N-terminal His-Thioredoxin tag in *E. coli* cultured in ZYM auto-inducible medium for 6 h at 37 °C, then overnight at 25 °C. Bacterial cells were then lysed by sonication, and cell debris and insoluble materials were removed by centrifugation. The supernatant was loaded into a Hitrap column, equilibrated with PBS 2× (buffer A) and 5 % of 100 mM Tris, pH 8.0, 300 mM imidazole buffer (buffer B). The protein was eluted by increasing the buffer B to 100 % over 20 mL. After dialysis overnight against 20 mM Tris, pH 8, 150 mM NaCl buffer, the His-Thioredoxin tag was cleaved off from the rest of the construct using HRV 3C protease. The protein was further purified by size-exclusion chromatography using a 26/60 Superdex 75 column equilibrated with 20 mM Tris, pH 8, 150 mM NaCl buffer. The protein was concentrated, flash-frozen in liquid N_2_, and stored at -80 °C. The protein concentration was determined using a BCA assay.

### Production and purification of Ist2-3C-Bad from yeast

W303.1b/*GAL4* (*a*, *leu2-3*, *his3-11*, *trp1-1:TRP1-GAL10-GAL4*, *ura3-1*, *ade2-1*, *can r*, *cir +*) yeast strain was transformed by the lithium-acetate method. Yeast cultures and recombinant protein expression were performed as previously described (66). Yeast cells were then harvested by centrifugation, washed with ice-cold deionized H_2_O, and then with ice-cold TEKS buffer (50 mM Tris-HCl pH 7.5, 1 mM EDTA, 0.1 M KCl, 0.6 M sorbitol), and resuspended in TES buffer (50 mM Tris-HCl, pH 7.5, 1 mM EDTA, 0.6 M sorbitol) supplemented with protease inhibitors (SIGMAFAST EDTA-free protease inhibitor cocktail) and 1 mM PMSF. The cells were subsequently broken with 0.5 mm glass beads using a ‘Pulverisette 6’ planetary mill (Fritsch). The crude extract was spun down at 1,000 × *g* for 20 min at 4 °C to remove cell debris and nuclei. The resulting supernatant was centrifuged at 20,000 × *g* for 20 min at 4 °C, yielding S2 supernatant and P2 pellet. The S2 supernatant was then ultracentrifuged at 125,000 × *g* for 1 h at 4 °C. The resulting P2 and P3 pellets were resuspended at 30–50 mg.mL^-1^ of total protein in TES buffer.

To purify Ist2 and Ist2ΔC, membranes obtained after expression of Ist2 (P2 or P3) were diluted to 5 mg.mL^-1^ of total protein in ice-cold buffer A (50 mM MOPS-Tris, pH 7.0, 500 mM NaCl, 20% (w/v) glycerol), supplemented with 1 mM PMSF and protease inhibitors. n-dodecyl-β-D-maltoside (DDM) was then added to 15 mg.mL^-1^, resulting in a DDM/protein ratio of 3/1 (w/w). The suspension was then stirred gently on a wheel for 1 h at 4 °C. Insoluble material was pelleted by centrifugation at 100,000 × *g* for 1 h at 4 °C. The supernatant was applied onto streptavidin-sepharose resin (1 mL *per* 3 mg Ist2) and incubated for 2 h at 6 °C to allow binding of BAD-tagged Ist2 to the resin. The resin was washed four times with three resin volumes of ice-cold buffer A supplemented with 0.5 mg.mL^-1^ DDM. Elution was performed by adding 70 µg of purified HRV-3C protease per mL of resin and overnight incubation at 6 °C. Before reconstitution, the eluted fraction was concentrated to 0.3–0.4 mg.mL^-1^ using a Vivaspin unit (100 kDa MWCO).

### Peptides

Ist2[729-768] peptide (SYGVAGATLPETIPTSKNYYLRFDEDGKSIRDAKSSAESS) and its scrambled version (DKYASNSAKSTYGIRIPVATFESLLRSKSGETDAYGEDPS) were from Proteogenix. Peptide purity was > 95%.

### Lipids

18:1/18:1-PC (1,2-dioleoyl-*sn*-glycero-3-phosphocholine or DOPC), 16:0/18:1-PC (1-palmitoyl-2-oleoyl-*sn*-glycero-3-phosphocholine or POPC), 16:0/18:1-PS (1-palmitoyl-2-oleoyl-*sn*-glycero-3 phospho-L-serine or POPS), brain PI(4)P (L-α-phosphatidylinositol 4-phosphate), brain PI(4,5)P_2_ (L-α-phosphatidylinositol 4,5-bisphosphate), NBD-PE (1,2-dioleoyl-*sn*-glycero-3-phosphoethanolamine-N-(7-nitro-2-1,3-benzoxadiazol-4-yl)), Rhodamine-PE (1,2-dipalmitoyl-*sn*-glycero-3-phosphoethanolamine-N-(lissamine rhodamine B sulfonyl), 16:0/12:0 NBD-PS (1-palmitoyl-2-(12-[(7-nitro-2-1,3-benzoxadiazol-4-yl)amino]dodecanoyl)-*sn*-glycero-3-phosphoserine) and 18:1 MPB-PE (1,2-dioleoyl-sn-glycero-3-phosphoethanolamine-N-[4-(p-maleimidophenyl)butyramide]) were purchased from Avanti Polar Lipids.

### Liposomes preparation

In glass tubes, lipids stored in CHCl_3_ or CHCl_3_/methanol stock solutions were mixed at the desired molar ratio. The tubes were pre-warmed at 33 °C for 5 min; then, the solvent was dried under a nitrogen flux for 25 min, and the tubes were placed in a vacuum chamber for 40 min to remove the remaining solvent. The lipid film was hydrated in 50 mM HEPES, pH 7.4, 120 mM K-Acetate (HK) buffer or 50 mM HEPES, pH 7.4, 210 mM sucrose buffer to obtain a suspension of multilamellar vesicles. The multilamellar vesicle suspensions were frozen and thawed five times and then extruded through polycarbonate filters of 0.2 µm pore size using a mini-extruder (Avanti Polar Lipids). Liposomes were stored at 4 °C and in the dark when containing fluorescent lipids and used within 2 days.

### Proteoliposome preparation

Liposomes were formed from DOPC or a POPC/POPS (9:1, mol/mol) mixture. Lipids were dissolved at 10 mg.mL^-1^ in 5 mL CHCl_3_ and dried in a rotavapor for approximately 30 min under pressure to form a thin lipid film in a glass balloon. After CHCl_3_ evaporation, the lipid film was placed under vacuum in a desiccator for at least 1 h to eliminate CHCl_3_ traces. Lipids were then suspended in 12.5 mL of buffer R (50 mM MOPS-Tris, pH 7, 200 mM NaCl), yielding multilamellar vesicles at a final 4 mg.mL^-1^ concentration. The vesicles were aliquoted, flash-frozen in liquid nitrogen, and stored at -80 °C. Immediately before reconstitution, 400 μL of the vesicles were thawed and allowed to equilibrate at room temperature. All subsequent steps were performed at room temperature. The vesicles were then solubilized for 15 min with Triton X-100 (TX-100) under agitation using a magnetic stirrer. The detergent: lipid ratio used was 2.5:1 (w/w) as determined by (67), resulting in final concentrations of 7 mg TX-100.mL^-1^ and 2.7 mg lipids.mL^-1^. Then, 2 mol% of NBD-PS resuspended in 0.5 mg.mL^-1^ DDM was added to the lipid/TX-100 mixture, along with 1 mM EGTA. The purified protein was added to the NBD-labelled detergent/lipid suspension at 14.4 mg.mmol^-1^ protein/lipid ratio (∼ 0.05 mg.mL^-1^ protein) and incubated for 30 min at room temperature. Triton X-100 and DDM were removed by adding pre-washed Bio-beads SM-2 adsorbent (Bio-Rad) in three steps: first, the sample was incubated for 2 h with 10 mg Bio-beads/mg TX-100, then a second aliquot of Bio-beads was added (10 mg/mg TX-100), and the sample was incubated for another 2 h. Lastly, Bio-beads from the first two steps were removed, and an additional 20 mg Bio-beads/mg was added for 1 h to the sample (67). Finally, the proteoliposomes were separated from the beads and either stored at 4°C for up to one week or flash-frozen in liquid N_2_ and stored at -80 °C.

### Preparation of the apo- and lipid-bound form of Osh6

For some experiments, Osh6 and its AF647-labelled version were prepared in an apo form or in a 1:1 complex with PS or PI(4)P. Protein (3.6 µM) was incubated with “heavy” liposomes (800 µM) composed of DOPC/POPS (95:5) or DOPC/brain PI(4)P (95:5), respectively, encapsulating 50□mM HEPES, pH 7.4, 210□mM sucrose buffer, in a volume of 250□μL of HK buffer. The apo form of Osh6 was prepared by incubating the protein with pure DOPC liposomes. Each sample was mixed with liposomes by agitation for 30□min at 30□°C and then centrifuged at 400,000 × *g* for 20□min at 20□°C to pellet the liposomes using a fixed-angle rotor (Beckmann TLA 120.1). A fraction of each supernatant (200□μL) containing Osh6 loaded with lipid was collected, and the concentration of each complex was assessed by measuring sample absorbance at λ = 280 nm (ε = 55,810 M^-1^.cm^-1^) and in the case of AF647-Osh6, at λ = 647 nm (ε = 265,000 M^-1^.cm^-1^).

### GST pull-down assay

A volume of 50 µL Glutathione Sepharose 4B beads (slurry) was combined with either GST, GST-Ist2[590–946], GST-Ist2[590–768], GST-Ist2[727–776] or GST-Ist2[590–946]Δ727–749 (2.5 µM final concentration) in a total volume of 150 µL of buffer [50 mM Tris, pH 7.4, 120 mM NaCl, 1 mM MgCl_2_, 1% Triton X-100, 1 mM DTT, 10% glycerol, and 0.25 mM PMSF]. The samples were incubated for 30 min at 4 °C under constant shaking (800 rpm). After washing the beads to remove unbound GST construct, Osh6 or its mutated forms, Osh4, ORD^Osh3^, or ORD^ORP8^ was added at a concentration of 2.5 µM in a final volume of 150 µL buffer. Samples were incubated at 25 °C for 1 h under constant shaking. Finally, the beads were washed, and the amount of bound proteins was analyzed on SDS-PAGE.

### Protein size measurement

The experiments were performed at 25 °C using a Wyatt DynaPro-99-E-50 System (Protein Solutions). A small volume of each purified protein (200 µL) was dialyzed using a Slide-A-Lyzer MINI dialysis device (MWCO 3.5 kDa) three times against HK buffer for 30 min to remove glycerol from protein stocks and to exchange buffer. Then, the samples were subjected to ultracentrifugation at 100,000 × *g* for 20 min at 20 °C to pellet the protein aggregate. A volume of 20 µL of the supernatant was added to the DLS quartz cuvette. A set of 12 autocorrelation curves was acquired, and the data were analyzed using the regularization algorithm of the Dynamics v6.1 software.

### Circular dichroism

The experiments were performed on a Jasco J-815 spectrometer at room temperature with a quartz cell of 0.05 cm path length (Starna Scientific Ltd.). Each protein was dialyzed in a Slide-A-Lyzer MINI dialysis device (MWCO 3.5 kDa) three times against 20 mM Tris, pH 7.4, 120 mM NaF buffer for 30 min to remove glycerol or DTT from the protein stock and to exchange buffer. Then, the samples were subjected to ultracentrifugation at 100,000 × *g* for 20 min at 20 °C to pellet protein aggregates, and the supernatant was collected. Each CD spectrum is the average of ten scans recorded from λ = 190 to 260 nm with a bandwidth of 1 nm, a step size of 0.5 nm, and a scan speed of 50 nm.min^-1^. Protein concentration was determined at λ = 280 nm by spectrophotometry or by densitometry after SDS-PAGE against a BSA concentration range. A control spectrum of buffer was subtracted from each protein spectrum. The percentages of protein secondary structure were estimated by analyzing their CD spectrum (in the 190– 250 nm range) using the BeStSel method provided online (68).

### Microscale thermophoresis

Ist2[729–768] peptide and its scrambled version were solubilized in 10 mM HEPES, pH 7.4, 150 mM NaCl buffer supplemented with 0.5 % Tween 20 and 3 mM EDTA (HBS-EP+ buffer) to prepare a stock solution. Ist2[590–936] was dialyzed in a Slide-A-Lyzer MINI dialysis device (MWCO 3.5 kDa) three times against HBS-EP+ buffer for 30 min to remove glycerol and DTT from protein stocks and to exchange buffer. The concentration of peptides and Ist2[590–936] solutions was determined by spectrophotometry at λ = 280 nm. Then a series of 16 1:1 dilution was prepared using HBS-EP+ buffer in a non-binding 96-well black plate (Greiner Bio-One), producing ligand concentrations ranging from 10^-9^ to 10^-3^ M. Each ligand dilution (50 µL) was mixed with 10 µL of a stock solution of Osh6-AF647 (120 nM), which led to a final concentration of Osh6-AF647 of 20 nM. The plate was incubated for 10 min under constant shaking at 25°C. Each sample was loaded in Premium Monolith NT.115 Capillaries (NanoTemper Technologies). MST was measured using a Monolith NT.115 instrument (NanoTemper Technologies) at an ambient temperature of 25°C. Instrument parameters were adjusted to 50 % LED power and high MST power. Data from three independent measurements were fitted with a non-linear regression model in GraphPad Prism using the signal from an MST-on time of 5 s to obtain a dissociation constant K_D_.

### CPM accessibility assay

The day of the experiment, 100 µL from a stock solution of Osh6(noC/S190C) construct was applied onto a 0.5 mL Zeba spin desalting column (7 kDa molecular weight cut-off) equilibrated with freshly degassed HK buffer, according to the manufacturer’s indications, to remove DTT. The concentration of the eluted protein was determined by spectrophotometry, considering ε = 55,810 M^-1^. cm^-1^ at λ = 280 nm. A stock solution of CPM (7-Diethylamino-3-(4-maleimidophenyl)-4-methylcoumarin, Sigma-Aldrich) at 4 mg/mL was freshly prepared as described in (69) by mixing 1 mg of CPM powder in 250 µL of DMSO. Thereafter, 50 µL of this solution was diluted in a final volume of 2 mL of HK buffer and incubated for 5 min at room temperature. The solution was protected from light and used immediately. In individual wells of a 96-well black plate (Greiner Bio-one), Osh6(noC/S190C) at 400 nM was incubated either with liposomes (400 µM total lipid) only made of DOPC or additionally containing 5% brain PI(4)P or POPS in 200 µL of HK buffer, in the presence or absence of 40 µM Ist2[729–768] peptide for 10 min at 25°C under constant shaking. Then, a small volume of CPM stock solution was added to obtain a final concentration of 4 µM. After a 25 min-incubation, fluorescence emission was measured at 465 nm (bandwidth 5 nm) upon excitation at λ = 387 nm (bandwidth 5 nm) using a fluorescence plate reader (TECAN M1000 Pro). Control intensities were recorded in the absence of protein for each condition.

### Flotation assay

Flotation assays using liposomes were performed as described in (70) with minor modifications. The association of Ist2 constructs with membranes was measured by mixing the protein (0.75 µM) with liposomes of the desired composition, doped with 0.1 mol% NBD-PE (750 µM lipids) for 1 h at 25 °C under constant shaking (800 rpm). For some experiments, this mixture was supplemented with 2 mM DTT before being mixed with Osh6 (0.75 µM) for 10 min under constant shaking. Next, in all cases, the liposome/protein mixture (final volume 150 µL) was adjusted to 28% (w/w) sucrose by mixing 100 µL of a 60% (w/w) sucrose solution in HK buffer and overlaid with 200 µL of HK buffer containing 24% (w/w) sucrose and 50 µL of sucrose-free HK buffer. The sample was centrifuged at 201,600 × *g* (average centrifuge force) in a swing rotor (TLS 55 Beckmann) for 70 min. The bottom (250 µL), middle (140 µL), and top (110 µL) fractions were collected. The bottom and top fractions were analyzed by SDS-PAGE after staining with SYPRO^®^ Orange using a FUSION FX fluorescence imaging system.

### Aggregation assays

The experiments were performed at 25 °C using a Wyatt DynaPro-99-E-50 System (Protein Solutions). L_A_ liposomes (50 µM total lipids) composed of DOPC/MPB-PE/NBD-PE (89.9:10:0.1) or DOPC/NBD-PE (99.9:0.1) were mixed with an equivalent amount of L_B_ liposomes composed of POPC/POPS/PI(4,5)P_2_/NBD-PE (64.9:30:5:0.1) or POPC/NBD-PE (99.9/0.1) in 20 µL of freshly degassed HK buffer and added to the quartz cell. A first set of 12 autocorrelation curves was acquired to measure the size distribution of the initial liposome suspension. Then, the cIst2[590–946] construct (500 nM final concentration) was added manually and mixed thoroughly. For all the experiments, aggregation kinetics were measured by acquiring one autocorrelation curve every 10 s for 80 min. At the end of the experiment, a set of 12 autocorrelation functions was acquired. The data were analyzed using two different algorithms provided by the Dynamics v6.1 software (Protein Solutions). The autocorrelation functions were fitted during the kinetics, assuming the size distribution is a simple Gaussian function. This mode, called the monomodal or cumulant algorithm, gives a mean hydrodynamic radius, R_H_, and width (or polydispersity). The polydispersity is represented in the kinetics measurements by the shaded area. It can reach tremendous values because of the simultaneous presence of free liposomes and liposome aggregates of various sizes. The autocorrelation functions were fitted before and after the aggregation process using a regularization algorithm that can resolve several populations of different sizes, such as free liposomes and liposome aggregates.

### Confocal microscopy with liposomes

Giant liposomes were produced by polyvinyl alcohol (PVA)-assisted swelling (71). Fifty microliters of 5% (w/w) PVA solution (average molecular weight 146,000–186,000, Sigma-Aldrich) was spread on a glass coverslip (diameter 1.5 mm) and dried at 55□°C for 30□min in an oven. Then, a volume of 20 µL of lipid mixture (1□mg/mL) with the desired lipid ratio in chloroform was spread on the top of the PVA layer. The solvent was allowed to evaporate overnight in a vacuum chamber. A volume of 200 µL of swelling buffer (20□mM HK, pH 7.5, 210□mM sucrose) was added to the coverslip to form giant liposomes. After an incubation of 20 min at room temperature, vesicles ( ∼ 130 µM total lipids) were collected by pipetting and kept at room temperature, protected from light. A volume of 10 µL of liposome suspension (2 mM lipids) was gently added to 50 µL of the suspension of giant liposomes. Then, a volume of 3.5 µL of a stock solution of DTT-free _C_Ist2[590–946] (28 µM) was gently added. After 30 min, the liposomes were imaged in µ-Slide 18 well Ibitreat (ibidi); wells were previously coated for 1□h with 2□mg/mL BSA and washed three times with degassed HK buffer. A volume of 5 µL of the mixture was added to 100 µL of HK buffer in each well for imaging. For the experiments with AF647-Osh6, a volume of 0.5 µL of the fluorescent protein (stock solution 18 µM) was previously added to the well. After a few minutes, the samples were observed at room temperature using a Leica TCS SP8 STED 3X in confocal mode. Images were acquired through a 63 × /1.4 NA oil objective using the LAS X software and analyzed using ImageJ software.

### NBD-PS transfer assays

These assays were performed using a Jasco FP-8300 spectrofluorometer at 30 °C with Osh6 and ORD^ORP8^. For experiments with three liposomes, L_A_ liposomes (200 µM lipids) composed of DOPC/MPB-PE/NBD-PS (88:10:2) were mixed with an equivalent amount of L_B_ liposomes composed of POPC/POPS/PI(4,5)P_2_/Rhod-PE (63:30:5:2) and L_C_ liposomes composed of POPC/POPS (70:30) in HK buffer. Then, a small volume of DTT-free _C_Ist2[590–946] (500 nM final concentration) or HK buffer was mixed for 10 min with the liposomes. The capacity of free MPB-PE lipids to form covalent bonds with thiol-containing proteins was then neutralized by adding 2 mM DTT. Upon adding Osh6 or ORD^ORP8^ (200 nM), the specific transfer of NBD-PS to L_B_ liposomes was measured for 11 min by following FRET between NBD-PS and Rhod-PE (λ_em_ = 580 nm, bandwidth = 2.5 nm) on excitation at λ_ex_ = 460 nm (bandwidth = 2.5 nm). Similar experiments were performed with L_B_ liposomes composed of POPC/POPS/PI(4,5)P_2_ (65:30:5) and L_C_ liposomes composed of POPC/POPS/Rhod-PE (68:30:2). The fluorescence signal (F) measured over time was normalized to the averaged signal measured just before the injection of Osh6 (F_0_). The speed of NBD-PS transfer was quantified by determining the time necessary to achieve half of PS equilibration between L_A_ and L_B_ (or L_C_) liposomes and calculating a rate corresponding to 1/t_1/2_.

For measuring PS transfer from proteoliposomes to liposomes, an 8-µL aliquot of proteoliposomes composed of DOPC/NBD-PS (98:2), containing Ist2, Ist2ΔC, or protein-free (“mock”), was diluted in 570 µL of HK buffer in the cuvette thermostated at 25°C under constant stirring. The fluorescence of the NBD signal was identical in each condition. After 400 s of incubation, L_B_ liposomes (50 µM lipids) made of POPC/POPS/PI(4,5)P_2_ (65:30:5) were added. Five minutes later, Osh6(HH/AA) was injected, and the transfer of NBD-PS to L_B_ liposomes was measured by following FRET between NBD-PS and Rhod-PE (λ_em_ = 580 nm, bandwidth = 2.5 nm) on excitation at λ_ex_ = 460 nm (bandwidth = 2.5 nm). The average signal (F_0_) measured before adding Osh6 was subtracted from the F value measured over time after adding Osh6. We also estimated the amount of NBD-PS that could be transported if all the molecules of NBD-PS from both the inner and outer leaflets of the PLs were accessible, by recording reference kinetics with twice the amount of “mock” PLs. The averaged F signal measured at the end of the kinetics, from which F_0_ was subtracted, was used to estimate the percentage of NBD-PS transferred from PLs to L_B_ liposomes by Osh6 from the other kinetics.

### PI(4,5)P_2_ transfer assay

This assay was carried out using a Jasco FP-8300 spectrofluorometer. A suspension (540 µL) of L_A_ liposomes (200 µM total lipid) composed of DOPC/PI(4,5)P_2_/Rhod-PE (93:5:2) was mixed with 250 nM NBD-PH_PLCδ1_ in HK buffer at 30°C. The mixture was in a cylindrical quartz cuvette, continuously stirred with a small magnetic bar, and thermostated at 30 °C. After 2 min, 60 µL of a suspension of L_B_ liposomes (200 µM lipids), only composed of DOPC, were injected. Osh6 or its HH/AA mutant (250 nM) was injected two minutes later. The signal was followed by measuring the NBD fluorescence emission at λ = 530 nm (bandwidth 5 nm) upon excitation at λ = 460 nm (bandwidth 5 nm) with a time resolution of 1 s. The amount of PI(4,5)P_2_ transferred from L_A_ to L_B_ liposomes was determined considering that [PI(4,5)P_2_] = 2.5 × F_Norm_ with F_Norm_ = (F-F_0_)/(F_Eq_-F_0_). F corresponded to the data point recorded over time, F_0_ was the average signal measured before the addition of Osh6, and F_Eq_ was the average signal measured in the presence of L_A-Eq_ and L_B-Eq_ liposomes containing 2.5% PI(4,5)P_2_. At equilibrium, it is considered that one half of accessible PI(4,5)P_2_ molecules contained in the outer leaflet of L_A_ liposomes (i.e., corresponding to 5% of 0.5 × 200 µM total lipids) have been transferred into L_B_ liposomes.

### Scramblase assay

An 8-µL aliquot of either “mock” or Ist2-containing proteoliposomes was diluted to a final 600 µL HK buffer volume. Then, the fluorescence of NBD-PS was monitored over time in a quartz cuvette, using excitation at 460 nm (bandwidth 2.5 nm) and emission at 534 nm (bandwidth 2.5 nm). After 5 min, 6 µL of a stock solution of sodium dithionite (from a 1 M dithionite stock solution, stored at -20 °C) was added (final concentration 10 mM), and the quenching of the NBD signal was measured for 5 min. For the experiments with Osh6(HH/AA), 200 nM of the protein was added 260 s before adding dithionite. Kinetics have been analyzed and/or normalized using the following formula: (F − F_end_)/(F_start_ − F_end_) × 100, where F is the intensity measured over time, F_start_ is the intensity just before dithionite addition, and F_end_ is the intensity after TX-100 addition (near 0), which allows to determine the extent of NBD reduction by dithionite.

### Iodide collisional quenching

A volume of 10 μL of liposomes (mock) or Ist2-containing PLs was diluted in 50 mM MOPS-Tris, pH 7, 40 mM Na_2_S_2_O_3_ buffer, directly in a quartz cuvette. Na_2_S_2_O_3_ was used to stabilize iodide ions in the solution. NBD fluorescence was followed by setting excitation and emission wavelengths at 470 nm and 530 nm, respectively, with slits of 5 nm, on a Horiba Jobin Yvon Fluorolog fluorimeter. Initial fluorescence (F_0_) was recorded for about 100 s, then concentrations of KI ranging from 0 to 0.2 M were added. KCl was used to adjust the ionic strength to 0.2 M for each cuvette before adding samples and KI. The mean fluorescence (F) value recorded over 240–290 s after KI addition was taken and used to plot ΔF (F_0_–F). The data were analyzed according to a modified Stern-Volmer equation (72, 73): F_0_/ΔF = (1/fa . K[Q]) + (1/fa) where F_0_ is the fluorescence intensity in the absence of the quencher, ΔF is the fluorescence intensity in the presence of the quencher at concentration [Q] subtracted from F_0_, fa represents the fraction of fluorescence accessible to iodide ions and K is the Stern-Volmer quenching constant.

### Modelling

Models of Osh6:Ist2 IDR heterodimer were obtained with AlphaFold3 (49) using *S. cerevisiae* Osh6 and Ist2[590–946] sequences as input (Uniprot, Q02201, and P38250, respectively). The identification of inter-chain H-bonds in the different models was done using Pymol.

### Plasmid, strains, and growth conditions for yeast experiments

Plasmids used for the yeast experiments are listed in **Table S2**. The NEBuilder Hifi DNA Assembly Kit was used to generate the following plasmids: pOsh6-mCherry-E170A, pOsh6-mCherry-3M, pIst2 derivative plasmids ([1-877], [590-946], [1-877]PM, short tail, short tail +23aa, short tail +40aa, very long tail, BS-Osh6-ER, BS-Osh6-PM) according to the supplier’s instructions. A site-directed mutagenesis kit (Agilent) was used to generate the deletion of the 736–743 region from the pIst2 plasmid. Briefly, the plasmid was amplified using oligonucleotides excluding the 736–743 region. All constructs were verified by DNA sequencing.

Yeast strains are listed in **Table S3**. Yeasts were grown in rich medium (YPD) or in synthetic dextrose (SD) medium containing 2% (wt/vol) glucose and appropriate amino acid drop-out mix (MP Biomedicals). Yeast was transformed using the standard lithium acetate/polyethylene glycol procedure. SD medium was supplemented with 1 mM ethanolamine when growing *cho1*-deficient strains or *psd1*Δ*ist2*Δ. The *ist2*Δ *OSH6-GFP TCB1-TagRFP* strain was constructed by homologous recombination using the appropriate marker as indicated in the table.

### Live yeast cell imaging

Yeast cells were grown for 14–18 h in appropriate SD medium at 30°C. Cells were diluted and harvested by centrifugation in mid-logarithmic phase (OD_600_=0.6–0.8) and prepared for visualization on glass slides, except for time-course experiments in which cells were maintained in a microfluidics chamber (see below). Imaging was conducted at room temperature, using a Spinning disk SR system (Olympus), equipped with an oil immersion plan apochromat 60x objective NA 1.42, an sCMOS Fusion BT camera (Hamamatsu), and a spinning-disk confocal system CSU-W1 (Yokogawa) except for cellular PS transport assay (see below). BFP-, GFP-, and mCherry/TagRFP-tagged proteins were visualized with DAPI (447/50), GFP (525/50), and mCherry (593/40) filters, respectively. Cells were imaged in 5 to 10 z-sections separated by 0.25 μm. Images were acquired using CellSens software (Olympus) and processed with Fiji (ImageJ).

### Cellular PS transport assay

Transport of PS in *cho1Δ* cells was performed as described previously (8, 48). Briefly, 18:1 lyso-PS (1-oleoyl-2-hydroxy-*sn*-glycero-3-phospho-L-serine, Avanti Polar Lipids) was dried under argon and resuspended in SD medium to 54 μM lyso-PS and used within 1–3 h after preparation. The PS transport assay was carried out using a Microfluidic Perfusion Platform (ONIX) driven by the interface software ONIX-FG-SW (Millipore). Strains *cho1*Δ and *cho1*Δ *ist2^736–743^*^Δ^, transformed with pC2 -GFP and other plasmids, as indicated, were grown to OD_600_ = 0.6–0.8 and injected into a YO4C microfluidics chamber and maintained in a uniform focal plane. When imagining two strains simultaneously, we stained one strain with the vacuolar dye CMAC (Life Technologies) at 100 µM for 10 min and washed twice before mixing the two strains at a ratio of 1:1. Normal growth conditions were maintained by flowing cells with SD medium or SD medium containing lyso-PS at 3 psi. Cells were imaged every 2 min over 30–40 min, in five *z*-sections separated by 0.7 µm. Imaging was performed at room temperature, using an Axio Observer Z1 microscope (Zeiss), equipped with an oil immersion plan apochromat 100× objective NA 1.4, an sCMOS PRIME 95 camera (Photometrics), and a spinning-disk confocal system CSU-X1 (Yokogawa). GFP-tagged proteins and CMAC/BFP-tagged proteins were visualized with a GFP Filter 535AF45 and a DAPI Filter 450QM60, respectively. Images were acquired with MetaMorph 7 software (Molecular Devices).

### Yeast image analysis

Images were processed with Fiji (ImageJ). Quantification of Osh6-mCherry localization was performed manually from a stack of 10 *z-*sections. Distribution of C2_Lact_–GFP was analyzed as described previously (8, 48). Briefly, a 3D stack of selected central z-sections over the entire time-course was constructed manually. Quantification of peripheral and internal peaks was performed by profiling cell signal intensity across a transversal line drawn in each cell at the start of the time-course, the external limit of the cell (perimeter) was selected, and total cell fluorescence was measured. Intensities of the peaks were quantified and normalized relative to the total signal. The cell profiles (peripheral and internal peaks) were followed and quantified over the time of the experiment. Data were processed in Excel and plotted using GraphPad.

### Yeast kinetic assays

The *psd1*Δ*ist2*Δ strains transformed with plasmids expressing BFP-Ist2 variants were grown for 14–18 h at 30 °C in SD-His medium supplemented with 2 mM ethanolamine. Cells were adjusted to an OD_600_ = 0.1 and grown in technical duplicates, in the presence or absence of ethanolamine, in 96-well microplates using the Spark^®^ microplate reader (Tecan). For each kinetic cycle, with an interval of 10 min, the cultures were shaken for 9 min with an amplitude of 2 mm and a frequency of 150 rpm, followed by a 1-min settling period before measuring the absorbance at 600 nm. Kinetic measurements continued until all strains reached the plateau of the stationary phase.

### Statistical analysis

Statistical analyses were performed using Prism (GraphPad). p values <□0.05, <□0.01, and <□0.001 are identified with 1, 2, and 3 asterisks, respectively. Ns: p□≥□0.05. The number of replicates (n) used for calculating statistics is specified in the figure legends.

## Supporting information

H-bonds between Osh6 and Ist2 residues identified in the five models of Osh6:Ist2 complex generated by AlphaFold3

## Acknowledgments

AF was supported by a fellowship from the Ministère de l’Enseignement Supérieur, de la Recherche et de l’Innovation. NA, CS, and this work were supported by an ANR-20-CE13-0030-02 and ANR-23-CE44-0026 grants from the Agence Nationale de la Recherche. We wish to thank Christine Jaxel for the full-length and ΔC pYeDP60-Ist2-3C-Bad constructs, Martin Cohen-Gonsaud and Laurène Heriaud for providing us with ORD^Osh3^, and Véronique Albanèse for the construction of the strain *ist2*Δ *OSH6-GFP TCB1-TagRFP.* We thank the joint IGMM-CRBM “yeast media and technologies service” and the MRI imaging facility, a member of the national infrastructure France-BioImaging, supported by the French National Research Agency (ANR-10-INBS-04, Investissements d’avenir).

## Data Availability

Raw data and source code are available from the corresponding author(s) upon reasonable request.

## Supplementary figures

**Figure S1.**
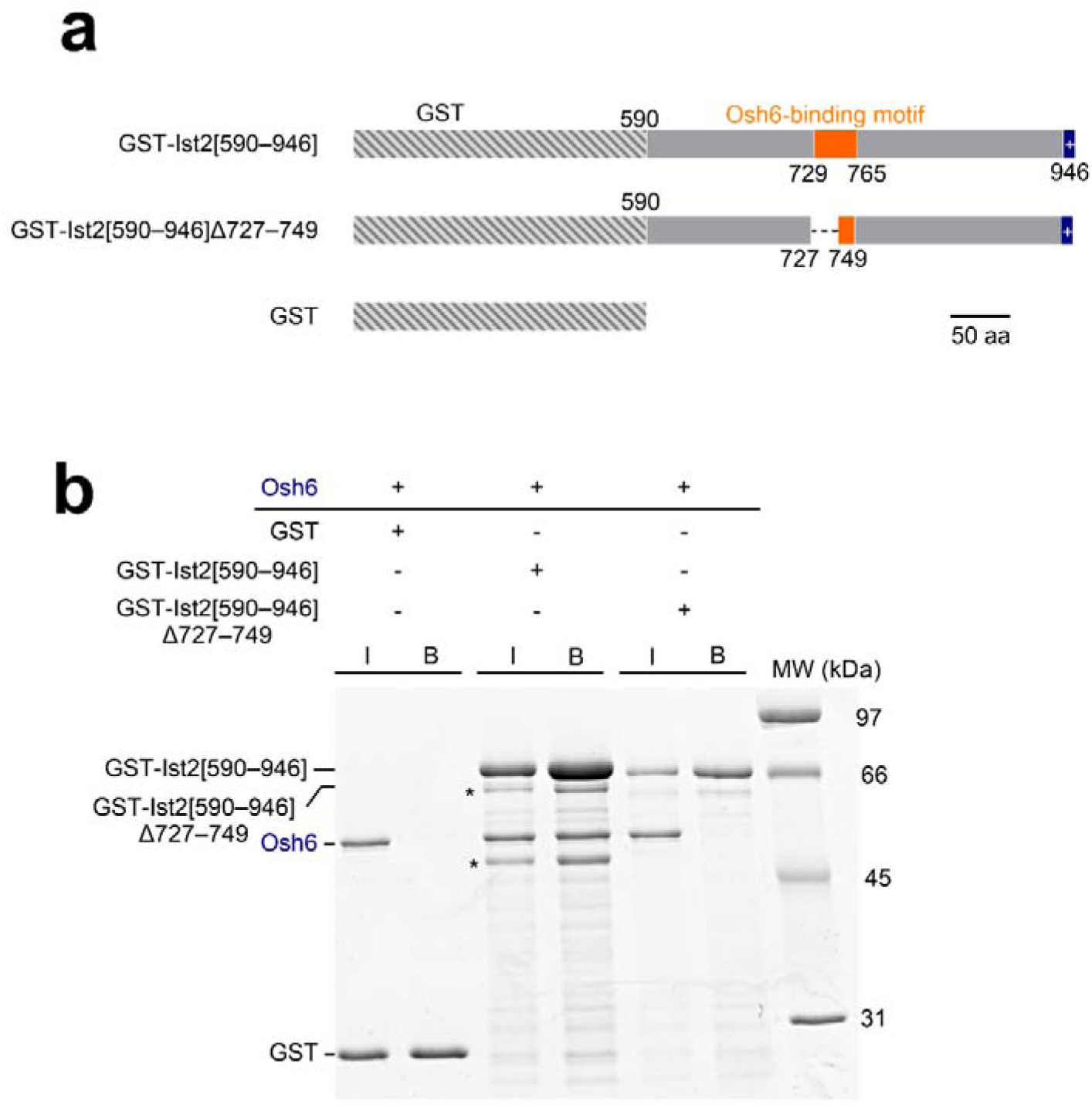
Additional GST pull-down assays with Osh6. Osh6 does not bind to the Ist2 IDR lacking a complete Osh6-binding motif. Purified GST, GST-Ist2[590– 946], and GST-Ist2[590–946]Δ727–749 were immobilized on glutathione beads and incubated with Osh6. The arrows indicate the different GST-Ist2 constructs, and the stars indicate contaminants.

**Figure S2.**
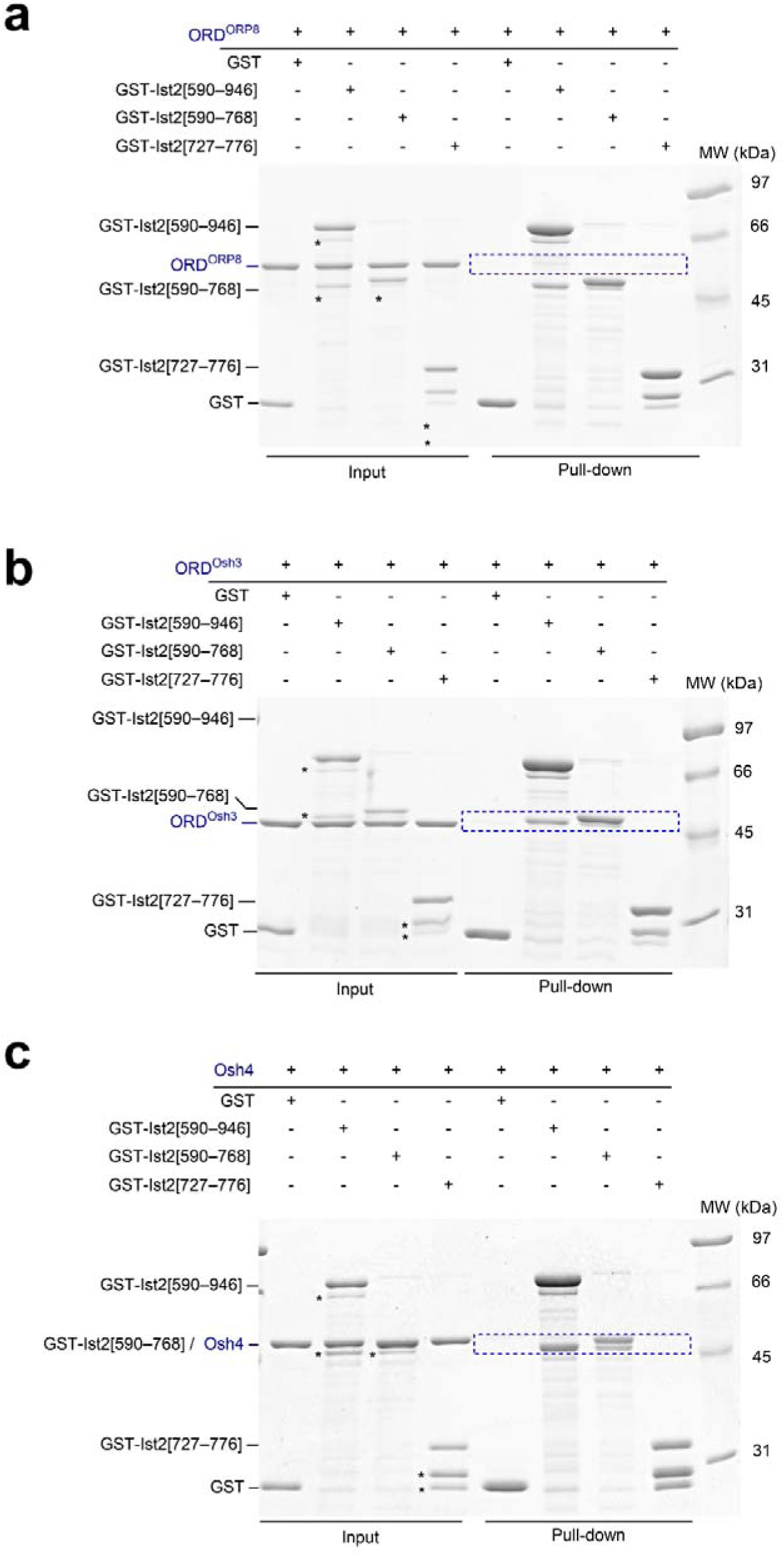
Additional GST pull-down assays with ORD^ORP8^, ORD^Osh3,^ and Osh4. Control experiments showed that **(a)** ORD^ORP8^, **(b)** ORD^Osh3,^ and **(c)** Osh4 do not bind to the Ist2 IDR. The experiments are performed as in Fig. 1b. Purified recombinant GST-Ist2[590–946], GST-Ist2[590–768], and GST-Ist2[727–776] constructs were immobilized on glutathione beads and incubated with ORD^ORP8^, ORD^Osh3,^ or Osh4. Input and bound fractions were analyzed using SDS-PAGE and SYPRO Orange staining. The arrows indicate the different GST-Ist2 constructs, and the stars indicate contaminants.

**Figure S3.**
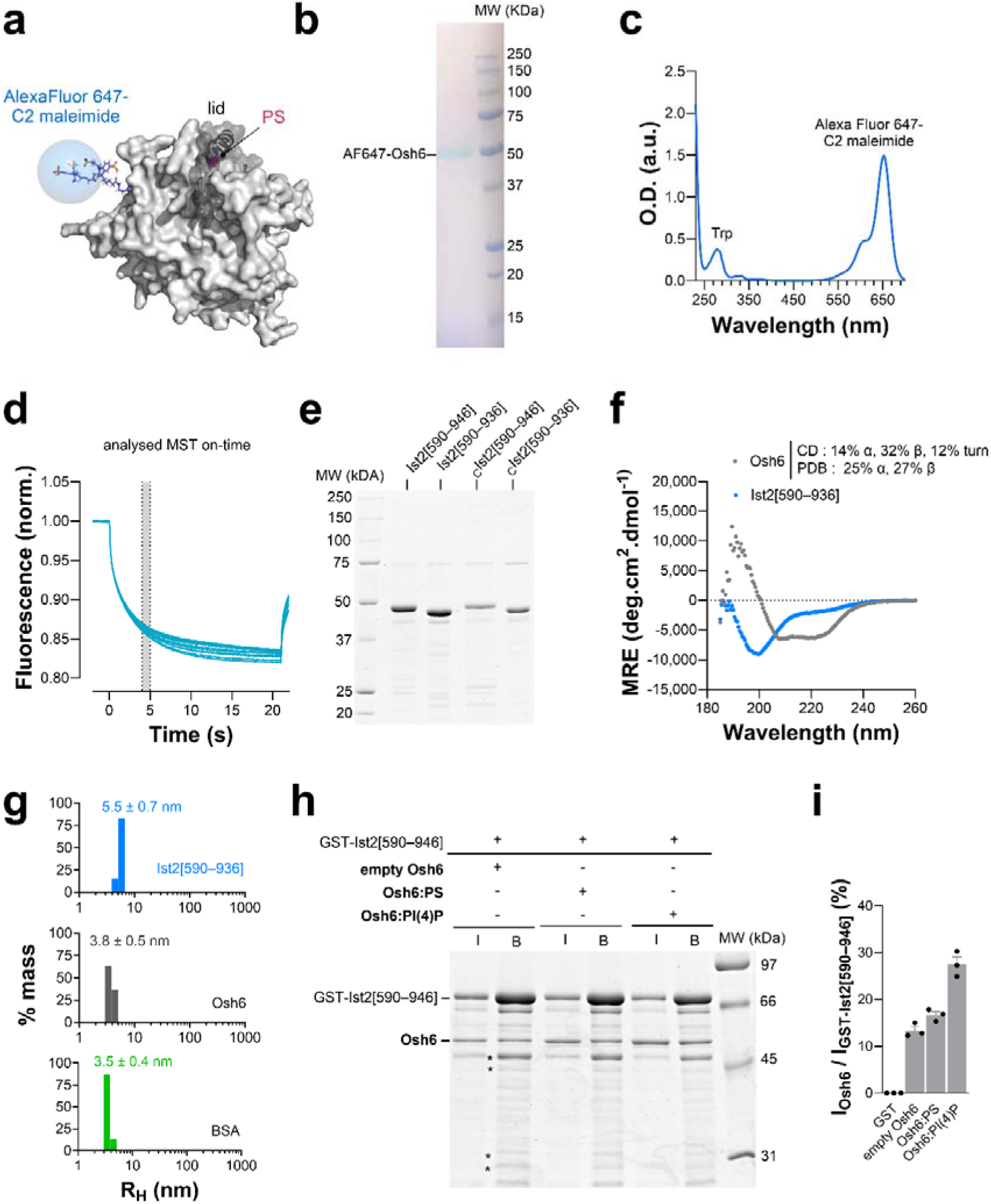
Measurement of Osh6:Ist2 binding affinity. **(a)** Three-dimensional model of AF647-Osh6 based on the crystal structure of Osh6 (PDB ID: 4B2Z). The solvent-exposed cysteines C62, C162, and C389 are mutated to serines; a cysteine replaces threonine T262. An Alexa Fluor 647-C2 maleimide moiety (in stick, with carbon in blue light, nitrogen in dark blue, and oxygen in red) is grafted to the thiol function of the C262 residue. The figure was prepared using PyMOL (http://pymol.org/). **(b)** SDS-PAGE of purified AF647-Osh6. The in-gel fluorescence was directly visualized without any staining. Lane 1: AF647-Osh6, lane 2: Precision Plus Protein™ All Blue Prestained Protein Standards (Bio-Rad). **(c)** UV-visible absorption spectrum of AF647-Osh6. Considering a purity grade of 100% for the protein, the optical density value at 280 nm (Trp) and 650 nm (Alexa Fluor 647-C2 maleimide) indicated that Osh6 was labelled with the dye in a 1:1 ratio. **(d)** Typical MST-based binding assay. The thermophoretic movement of AF647-Osh6 changes upon binding to Ist2[529–768] peptide. **(e)** SDS-PAGE of purified Ist2[590-936] and Ist2[590-946] constructs with or without an extra N-terminal cysteine. **(f)** Far-UV CD spectrum of purified Ist2[590–936] (17.7 µM) and Osh6 (12.8 µM) in 20 mM Tris, pH 7.4, 120 mM NaF buffer at room temperature. The percentage of α-helix, β-sheet, and turn, derived from the analysis of the spectrum of Osh6, are given, as well as the values derived from the crystal structure (PDB ID: 4B2Z) using the DSSP algorithm. MRE, mean residue ellipticity. **(g)** Hydrodynamic radius distribution by mass for Ist2[590–936] (35 µM; 38 kDa), Osh6 (50 µM, 51.6 kDa), and BSA (30 µM, 66.5 kDa) measured in HK buffer at 25 °C. **(h)** GST pull-down with Osh6 in an apo or lipid-bound state. Osh6 was mixed with liposomes made of DOPC or liposomes additionally containing 5 mol% POPS or brain PI(4)P, and separated from these liposomes by ultracentrifugation. Each form of the protein was incubated with GST-Ist2[590–946] immobilized on beads. Input (I) and bound (B) fractions were analyzed using SDS-PAGE. **(i)** Quantification of the GST-pull down assays as shown in (h).

**Figure S4.**
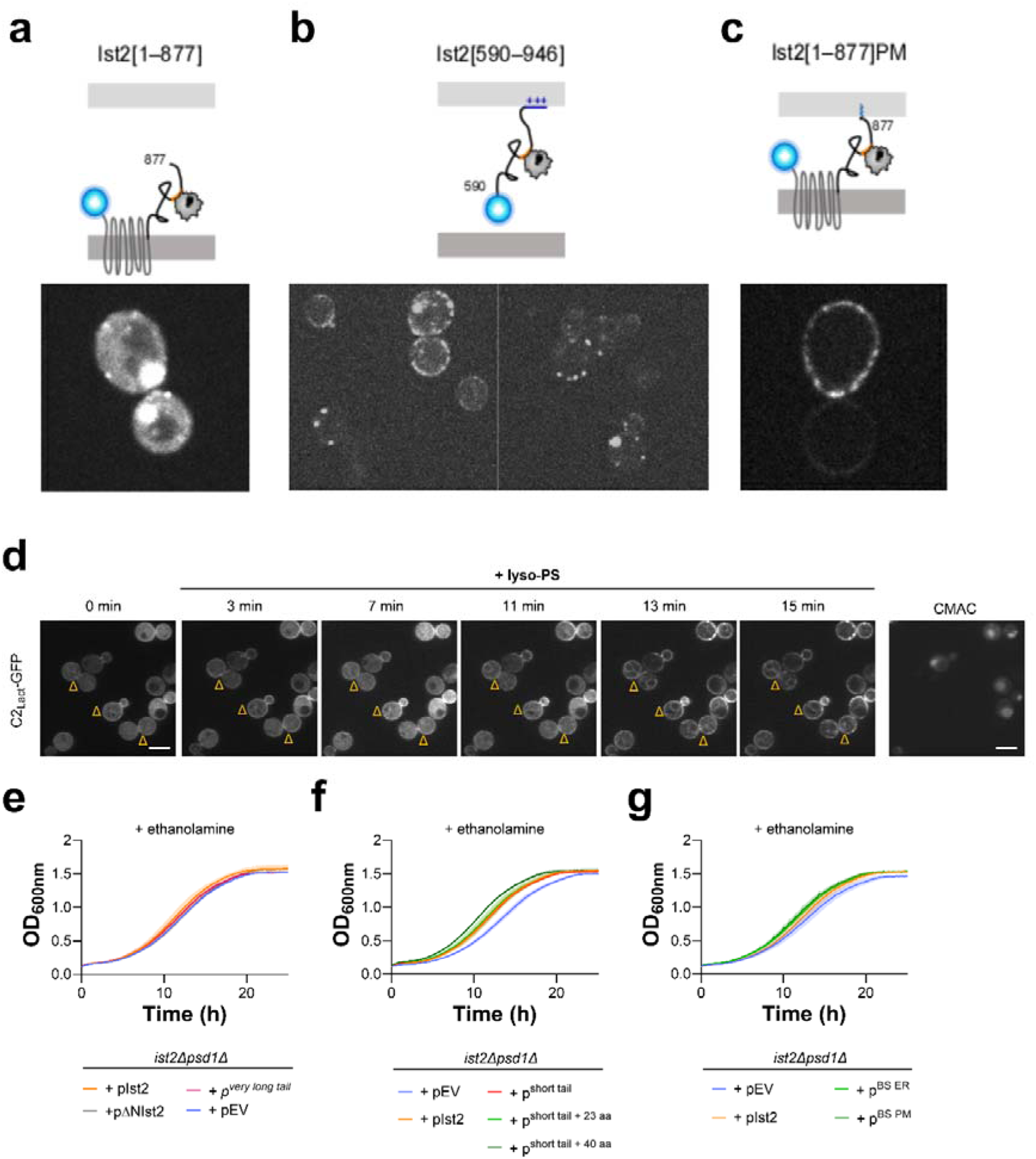
Supplementary information for Fig. 2: Localization and functionality of Ist2 truncation mutants in yeast. **(a)** Diagram of BFP-Ist2[1–877], lacking the PM binding region, and its localization in *ist2Δ* cells (lacking endogenous Ist2) (bottom panel). The bright spots likely represent BFP-Ist2[1–877] aggregates. This construct contains the full ER-embedded TM domain [1–589] and the cytosolic tail [590–878], with the Osh6-binding site depicted in orange, and Osh6 in grey. BFP is shown as a blue sphere. **(b)** Diagram of BFP-Ist2[590–946], lacking the full TM domain, and its localization in *ist2Δ* cells (bottom panels, showing two fields of cells). Note the low fluorescent signal; the puncti likely represent BFP-Ist2[590–946] aggregates. The PM-binding polybasic region is shown in blue, +++. **(c)** Diagram of BFP-Ist2[1–877]PM, where the PM binding region is replaced by the CAAX prenylation motif that allows attachment of the IDR to the PM via the lipid anchor (shown in green). Bottom panel shows that this construct localizes to the cortex of *ist2Δ* cells, and colocalizes with Osh6-GFP (not shown). **(d)** Snapshots of the cellular PS transport assay, showing the localization of the PS-probe C2_Lact_-GFP over time after the addition of lyso-PS. A mix of two strains was imaged in this experiment, *cho1*Δ and *cho1*Δ *ist2^736–743^*^Δ^ (marked with Δ in the images); the *cho1*Δ strain was stained with the vacuolar dye CMAC (right-most panel) before the experiment to distinguish between the two strains. Quantification of this experiment is shown in Fig. 2c. **(e-g)** Control growth curves of *ist2Δ psd1Δ* cells transformed with different pBFP-Ist2 plasmids, as indicated, where pEV represents an empty control plasmid. Yeast growth was monitored over time in minimal media in the presence of ethanolamine at 30°C.

**Figure S5.**
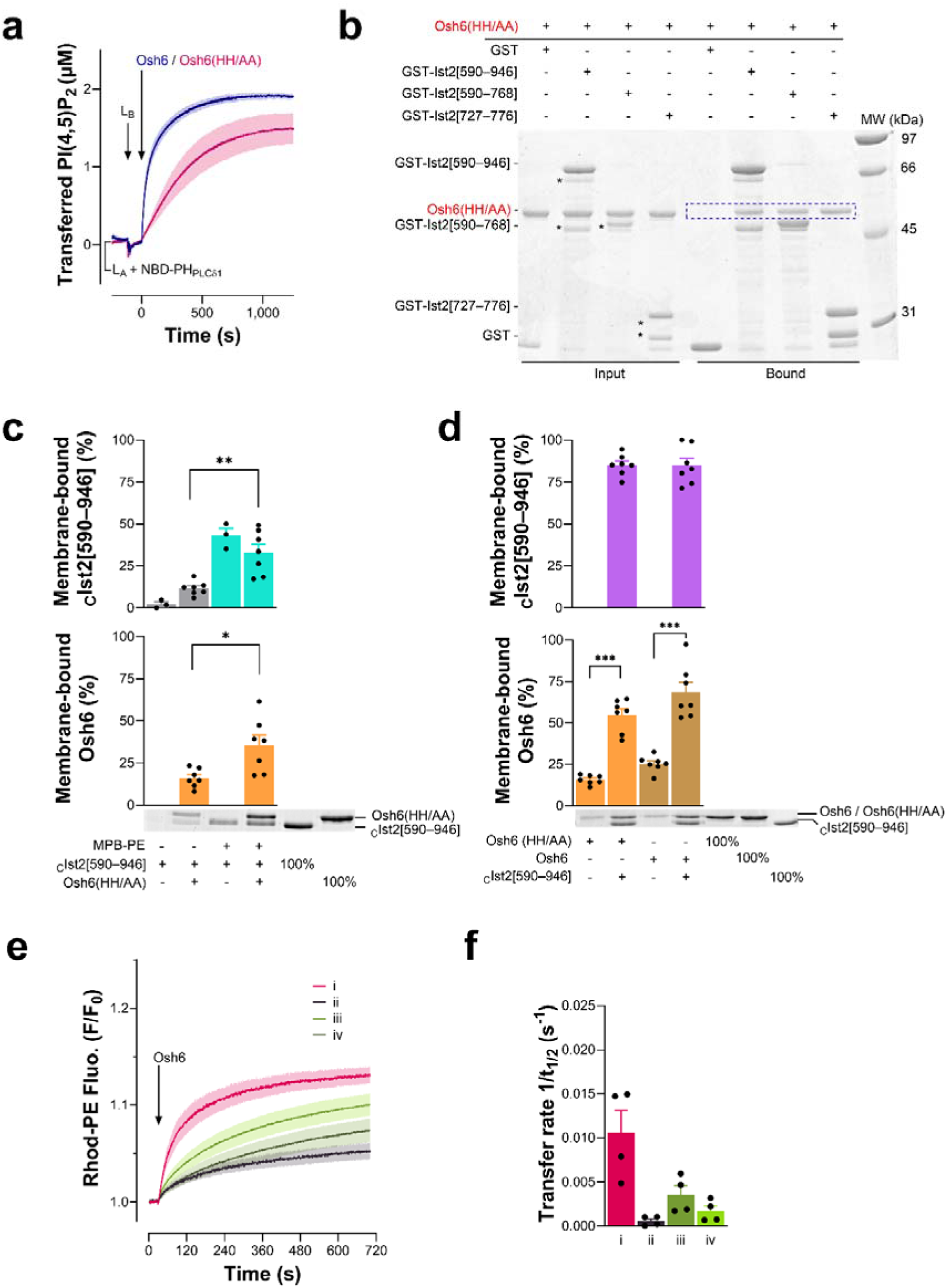
Lipid transfer activities of Osh6 and Osh6(HH/AA) and interaction with Ist2 IDR. **(a)** PI(4,5)P_2_ transfer activity of Osh6 and Osh6 (HH/AA). L_A_ liposomes (200 µM total lipids) composed of DOPC/PI(4,5)P_2_/Rhod-PE (93:5:2) were mixed with NBD-PH_PLCδ1_ (250 nM) at 30 °C. After 2 min, DOPC liposomes (200 µM, L_B_) were added. Two minutes later, Osh6 or Osh6(HH/AA) was injected (250 nM). The signal was normalized in terms of transferred PI(4,5)P_2_. Each curve is the mean ± s.e.m. of independent kinetics traces (n = 3). **(b)** GST pull-down. GST-tagged segments of the Ist2 IDR were immobilized on glutathione beads and incubated with Osh6(HH/AA). Input and bound fractions were analyzed on SDS-PAGE with SYPRO Orange staining. The stars indicate the main contaminants in GST-tagged construct preparation. As pointed out by the blue box, Osh6(HH/AA) interacts with the three GST-tagged segments of Ist2 IDR. No interaction was seen with GST alone. **(c)** Flotation assay. _C_Ist2[590–946] construct (0.75 µM) was mixed for 1 h with DOPC liposomes, containing or not 10% MPB-PE, and doped with 0.1% NBD-PE. Then, DTT (1 mM) was added to stop the functionalization reaction, and the liposomes were mixed or not with 0.75 µM of Osh6(HH/AA) for 10 min. The amount of membrane-bound cIst2[590–946] and Osh6(HH/AA) was determined using the content of lanes 5 and 6 (100% total), respectively. Data are represented as mean ± s.e.m. (n = 3–7) with single data points. Unpaired Mann–Whitney U test (\**p* < 0.05, \*\**p* < 0.01). **(d)** Flotation assay. Liposomes (750 µM lipids) composed of POPC/POPS/PI(4,5)P_2_/NBD-PE (65:30:5:0.1) were mixed with 0.75 µM of Osh6 or Osh6(HH/AA) in the presence or absence of an equivalent amount of cIst2[590–946] for 10 min. Data are represented as mean ± s.e.m. (n = 7) with single data points. Unpaired Mann–Whitney U test (\*\*\**p* < 0.001). **(e)** Real-time PS transport assays. (i) Osh6 was added to L_A_ liposomes (200 µM), doped with MPB-PE and 2% NBD-PS, and connected by _C_Ist2[590–946] (0.5 µM) to L_B_ liposomes (200 µM), containing PS, PI(4,5)P_2_ and 2% Rhod-PE in the presence of free L_C_ liposomes (200 µM) with no PI(4,5)P_2_. The increase in rhodamine fluorescence over time corresponds to NBD-PS transfer from L_A_ to L_B_ liposomes. A mirror experiment (ii), in which Rhod-PE is incorporated in L_C_ and not L_B_ liposomes, was conducted to measure the specific transfer of PS from L_A_ to L_C_ liposomes. These two experiments were also performed without _C_Ist2[590–946] (iii, iv). Each curve represents the mean ± s.e.m. of several kinetics recorded during independent experiments (n = 3). **(f)** Rate (1/*t*_1/2_) of NBD-PS transfer measured with Osh6 in conditions (i), (ii), (iii), and (iv).

**Figure S6.**
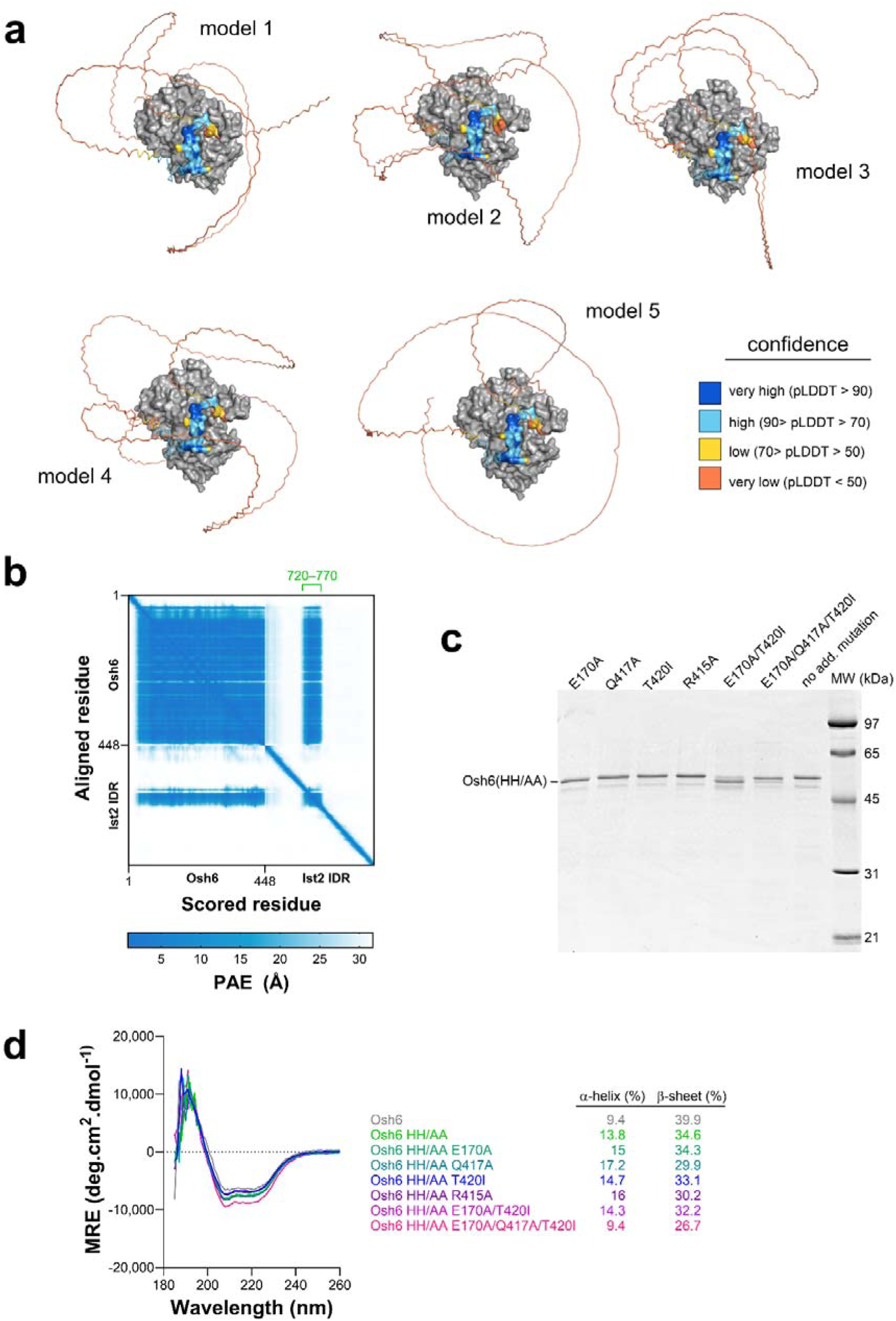
Identification of the Ist2-binding region on Osh6’s surface. **(a)** Five models of the Osh6:Ist2 IDR complex generated by AlphaFold3. Osh6 is represented as a grey surface. For clarity, the N-terminal low complexity sequence of Osh6 (35 residues) has been omitted. The Ist2 IDR is represented in ribbon mode and colored according to the predicted local distance difference test (pLDDT) *per* atom. The Ist2[729–747] segment (minimal Osh6-binding motif) is represented as a surface. **(b)** PAE is presented as a two-dimensional plot where the color at coordinates (x, y) represents the predicted position error at residue x if the expected and actual structures were aligned on residue y. The PAE values calculated for residue pairs from Osh6 and Ist2 IDR suggest that the position and orientation of the segment [720-770] of the Ist2 IDR relative to Osh6 is predicted with high confidence (lowest PAE values). **(c)** SDS-PAGE analysis of purified Osh6(HH/AA) constructs bearing additional mutations. **(d)** Far-UV CD spectrum of purified Osh6 and mutants (6.85–11.88 µM) in 20 mM Tris, pH 7.4, 120 mM NaF buffer at room temperature. The percentages of α-helix and β-sheet, derived from the analysis of each spectrum, are given, as well as the values derived from the crystal structure (PDB ID: 4B2Z) using the DSSP algorithm. MRE, mean residue ellipticity.

**Figure S7.**
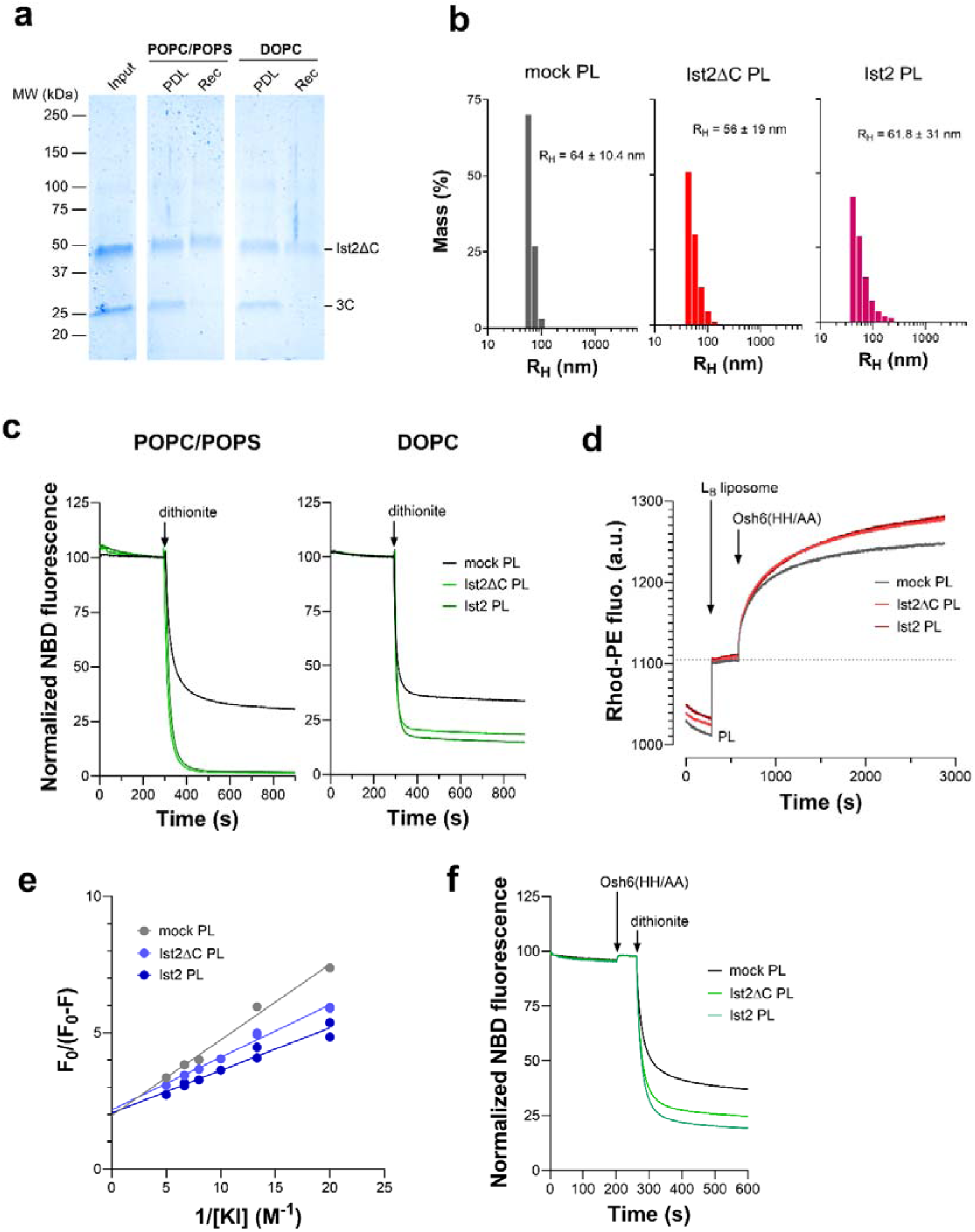
Osh6-mediated PS transfer is sustained by Ist2 PS scrambling activity. **(a)** Example of reconstitution. Ist2ΔC was reconstituted in liposomes composed of POPC/POPS/NBD-PS (89:9:2) or DOPC/NBD-PS (98:2) at ∼ 1/7000 protein/lipid molar ratio. Therefore, the protein/detergent/lipid (PDL) mixture contains 0.5 µM protein and 3.6 mM lipids. The reconstitution buffer was 50 mM MOPS-Tris, pH 7, and 200 mM NaCl. ‘Input’ corresponds to the protein sample after affinity purification on streptavidin beads. About 1 µg was loaded on the gel. PDL, protein/detergent/lipid mixture, before removal of the detergent. Rec, reconstituted samples after detergent removal with Biobeads. Samples were analyzed on a 4–15% TGX precast gel and were stained with Coomassie Blue. **(b)** Hydrodynamic radius distribution by mass of mock, Ist2, and Ist2ΔC proteoliposomes diluted in HK buffer at 25 °C. **(c)** Scramblase assay. NBD-PS-containing proteoliposomes, in which the Ist2ΔC or Ist2 construct is reconstituted, or protein-free liposomes (“mock”), were treated with dithionite (40 mM) to analyze the movement of NBD-PS between proteoliposome leaflets. Assays were done in 50 mM MOPS-Tris, pH 7, 200 mM NaCl, under stirring at 20 °C. The initial fluorescence was recorded for 300 s before the addition of dithionite to quench NBD-lipids on the outer leaflet. **(d)** Real-time PS transport assays. These raw kinetics correspond to the normalized curves shown in Fig. 7b. **(e)** Collisional quenching of the fluorescence of NBD-PS present in the membrane of mock and Ist2-containing PLs by iodide ions. The data are as modified Stern-Volmer plots (see Materials and Methods). F_0_ is the fluorescence intensity of the sample in the absence of quencher, whereas F is the fluorescence intensity at a given iodide ion concentration. The data points were fitted to a linear regression. The y-intercept of the linear regression for “mock” PLs is 1.962 ± 0.206, whereas it is 2.168 ± 0.107 for Ist2 PLs and 2.052 ± 0.157 for Ist2ΔC PLs. This means that ∼51 % of NBD-lipids in the membrane of “mock’ PLs are accessible to the quencher, whereas ∼46% and ∼49% of NBD-lipids in the membrane of Ist2-containing proteoliposomes are accessible to the quencher. **(f)** Scramblase assay. Osh6 (200 nM) was added to mock, Ist2ΔC, and full-length Ist2 PLs before dithionite treatment (10 mM).

**Table S1.**
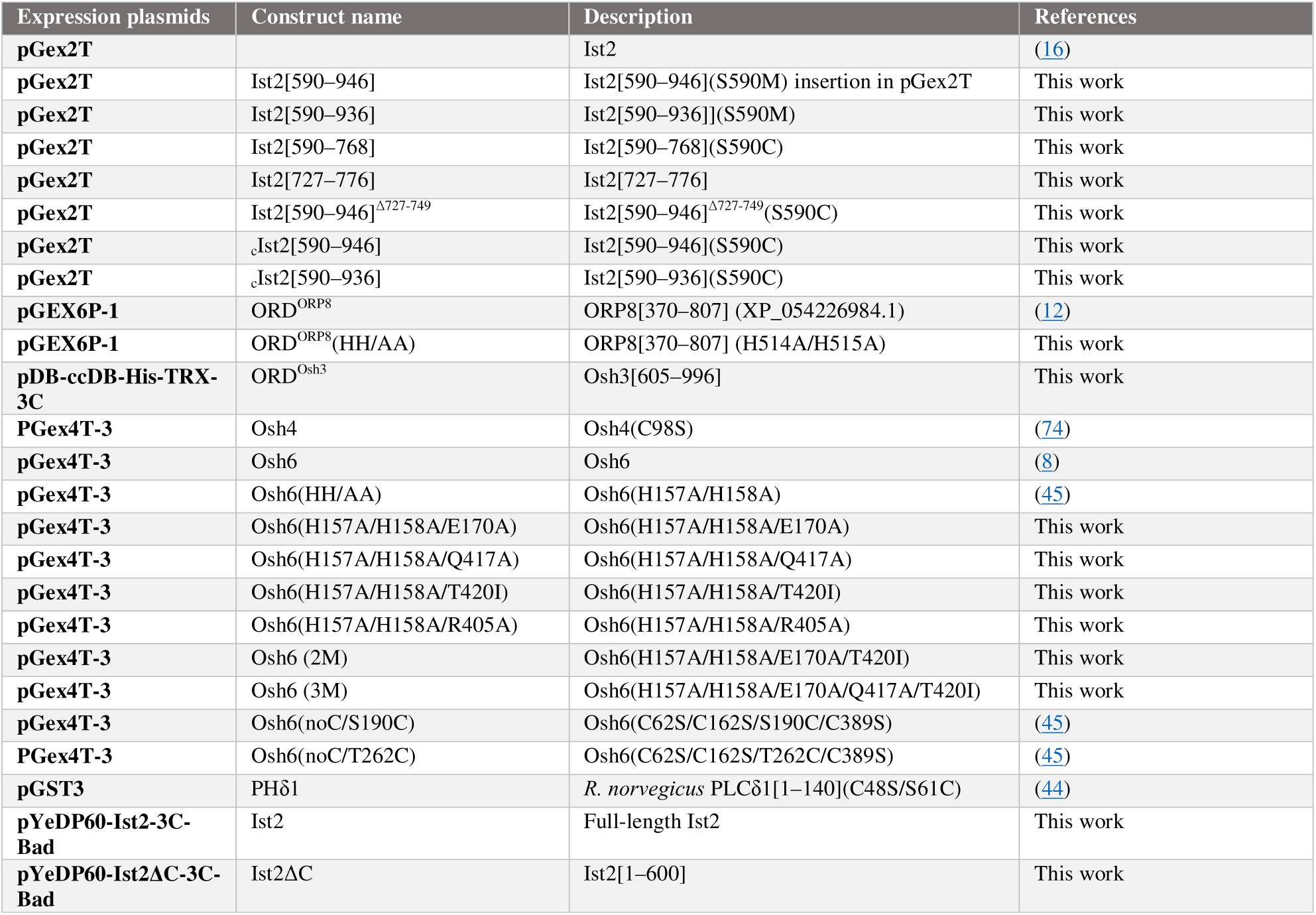
Plasmids and inserts used to produce recombinant proteins.

**Table S2:** Plasmids for yeast experiments used in this study. Note that all plasmids carry ampicillin antibiotic resistance. For mutated versions, the retained codon positions are indicated in brackets.

| Plasmids | Alias | Description | References |
| --- | --- | --- | --- |
| **pRS416** |  | *CEN, URA3* | Lab collection |
| **pOsh6-mCherry** | pAC100 | pRS416-based, ADH1pr-Osh6-mCherry | (16) |
| **pCS_39** | pOsh6-mCherry-E170A | ADH1pr-Osh6-mCherry, *E170A*; Mutations inserted by Gibson cloning from a pGex4T-3 plasmid expressing Osh6(E170A) into pAC100 plasmid | This work |
| **pCS_41** | pOsh6-mCherry-3M | ADH1pr-Osh6-mCherry, *E170A, Q417A, T420I*; Mutations inserted by Gibson cloning from a pGex4T-3 plasmid expressing Osh6(E170A/Q417A/T420I) into pAC100 plasmid | This work |
| **pRS413** | pRS413/pEV | *CEN, HIS3* | Lab collection |
| **pAK75** | GFP-Ist2 | pUG34-based (*CEN, HIS3*), Ist2pr-BFP-Ist2 | (22) |
| **pJMD_07** | pIst2 | pUG34-based (*CEN, HIS3*), Ist2pr-BFP-Ist2 | (16) |
| **pJMD_13** | pIst2[1–877] | Ist2pr-BFP-Ist2[1–877]; Exclusion of [878–946] region from pAK75 recombined into pJMD_07 | This work |
| **pJMD_10** | pIst2[590–946] | Ist2pr-BFP-Ist2[590–946]; Exclusion of [1–590] region from pAK75 recombined into *BamHI/XhoI* linearized pJMD_07 | This work |
| **pJMD_30** | pIst2[1–877]PM | Ist2pr-BFP-Ist2[1–877]-CAAX; insertion of CAAX-coding motif by Gibson cloning into pJMD_07 | This work |
| **pCS_45** | pIst2<sup>736-743Δ</sup> | Ist2pr-BFP-Ist2 [1–736; 743–946]; Site-directed mutagenesis of pJMD_07 | This work |
| **pJMD_20** | pIst2-short-tail | Ist2pr-BFP-Ist2-[1–590; 705–762; 879–946]; Synthetic gene integrated by Gibson cloning into pJMD_07 | This work |
| **pJMD_33** | pIst2-short-tail+23aa | Ist2pr-BFP-Ist2-[1–590; 682–762; 879–946]; Extension of 23 amino acids [682–704] by Gibson cloning into pJMD_20 | This work |
| **pJMD_34** | pIst2-short-tail+40aa | Ist2pr-BFP-Ist2-[1–590; 665–762; 879–946]; Extension of 40 amino acids [665–704] by Gibson cloning into pJMD_20 | This work |
| **pJMD_36** | pIst2-very-long-tail | Ist2pr-BFP-Ist2-[1–704; 590–704; 705–877; 763–946]; Duplication of [590–704] and [763–877] regions inserted by Gibson cloning into pJMD_20 | This work |
| **pJMD_18** | pIst2-BS-Osh6-ER | Ist2pr-BFP-Ist2-[1–589; 705–762; 590–704; 763–946]; Position change of [705–762] region inserted by Gibson cloning into pJMD_20 | This work |
| **pJMD_19** | pIst2-BS-Osh6-PM | Ist2pr-BFP-Ist2-[1–704; 763–877; 705–762; 878–946]; Position change of [705–762] region inserted by Gibson cloning into pJMD_20 | This work |
| **LactC2-GFP** | pC2lact-GFP | GPDprC2Lact-GFP, *CEN, URA3*, Addgene plasmid # 22852 | (75) |

**Table S3:** Yeast strains used in this study. Numbers after the Δ symbol indicate the starting position or the codon range of the deleted region

| *S.cerevisiae*<br>S288 Alias | Strains | Background | Genotype | References |
| --- | --- | --- | --- | --- |
| **SGAY1635** | *osh6Δosh7Δ* | SGA Y7039 | *MATa can1Δ::STE2pr-LEU2 lyp1Δ ura3Δ0 leu2Δ0 his3Δ1 met15Δ0 osh6Δ::HygMX osh7Δ::NatMX* | (7) |
| **MdPY01** | *ist2Δosh6Δosh7Δ* | SGA Y7039 | *MATa can1Δ::STE2pr-LEU2 lyp1Δ ura3Δ0 leu2Δ0 his3Δ1 met15Δ0 ist2Δ::KanMX osh6Δ::HygMX osh7Δ::NatMX* | (16) |
| **VAY2802** | *ist2Δ OSH6-GFP* | BY4742 | *MATa his3Δ1 leu2Δ0 lys2Δ0 ura3Δ0 ist2Δ::Hyg OSH6-GFP::KanMX* | (16) |
| **MdPY12** | *ist2Δ OSH6-GFP TCB1-TagRFP* | BY4742 | *MATa his3Δ1 leu2Δ0 lys2Δ0 ura3Δ0 ist2Δ::Hyg OSH6-GFP::KanMX TCB1-TagRFP::URA3* | This work |
| **MdPY08** | *cho1Δ* | BY4742 | *MATa his3Δ1 leu2Δ0 lys2Δ0 ura3Δ0 cho1Δ::kanMX* | (16) |
| **MdPY07** | *cho1Δ ist2<sup>736-743</sup>Δ* | BY4742 | *MATa his3Δ1 leu2Δ0 lys2Δ0 ura3Δ0 cho1Δ::kanMX ist2Δ736-743* | (16) |
| **MdPY11** | *psd1Δ ist2Δ* | BY4741 | *MATa his3Δ1 leu2Δ0 met15Δ0 ura3Δ0 psd1Δ::kanMX ist2Δ::natMX* | (28) |

## Supplementary File

Excel sheet listing the H-bonds between Osh6’s and Ist2’s residues identified in the five models of Osh6:Ist2 complex generated by AlphaFold3. A final table reports the number of times each H-bond is identified. The type of H-bonds is named as follows: sm, H-bond between the side-chain of an Osh6’s residue and the main chain of an Ist2’s residue; ms, m, H-bond between the side-chain of an Ist2’s residue and the main chain of an Osh6’s residue; ss, H-bond between the side-chain of an Osh6’s residue and the side-chain of an Ist2’s residue; mm, H-bond between the main chain of an Osh6’s residue and the main chain of an Ist2’s residue.

